# Universal features shaping organelle gene retention

**DOI:** 10.1101/2021.10.27.465964

**Authors:** Konstantinos Giannakis, Samuel J. Arrowsmith, Luke Richards, Sara Gasparini, Joanna M. Chustecki, Ellen C. Røyrvik, Iain G. Johnston

## Abstract

Mitochondria and plastids power complex life, and retain their own organelle DNA (oDNA) genomes, with highly reduced gene contents compared to their endosymbiont ancestors. Why some protein-coding genes are retained in oDNA and some lost remains a debated question. Here we harness over 15k oDNA sequences and over 300 whole genome sequences with tools from structural biology, bioinformatics, machine learning, and Bayesian model selection to reveal the properties of genes, and associated underlying mechanisms, that shape oDNA evolution. Striking symmetry exists between the two organelle types: gene retention patterns in both are predicted by the hydrophobicity of a protein product and its energetic centrality within its protein complex, with additional influences of nucleic acid and amino acid biochemistry. Remarkably, retention principles from one organelle type successfully and quantitatively predict retention in the other, supporting this universality; these principles also distinguish gene profiles in independent endosymbiotic relationships. The identification of these features shaping organelle gene retention both provides quantitative support for several existing evolutionary hypotheses, and suggests new biochemical and biophysical mechanisms influencing organelle genome evolution.

## Introduction

Mitochondria and plastids (the broader class of organelles of which chloroplasts are one type) are bioenergetic organelles derived from the ancient endosymbiotic acquisition of bacterial precursors [1]. The subsequent co-evolution of mitochondria and plastids with their host cells has shaped complex life [2, 3, 4]. Across eukaryotes, the genomes of the original endosymbionts (estimated to have contained thousands of genes [5]), have been dramatically reduced through evolutionary time [6, 7, 1]. Genes have either been lost completely or transferred to the ‘host’ cell nucleus, so that modern-day organelle DNA (oDNA) contains few genes, with profound implications for the balance of control between the nucleus and endosymbiont, and the inheritance and maintenance of vital genetic information [8].

Selective pressures favouring organelle gene transfer are largely agreed upon [7]. Nuclear encoding allows recombination to avoid Muller’s ratchet (the irreversible buildup of damaging mutations) [9, 6], protection from chemical mutagens [10, 11] and replication errors [12, 13], and enhanced fixing of useful mutations [7, 6]. However, these observations raise the dual question: why are any genes retained in organelles at all [14]? This question has been hotly debated over decades, with many proposed hypotheses. The preferential retention of genes encoding hydrophobic products has been suggested, due to the challenge of correctly targetting and importing such products to the correct organelle [15, 16, 17]. The retention of genes playing central roles in controlling redox activity has also been proposed, to facilitate local control of activity [18]. Other hypotheses, including roles for nucleic acid biochemistry [19], gene expression levels [20], energetic costs of encoding [21], toxicity [22], and others have been proposed, but quantitative testing of these ideas remains limited [19, 23].

Applying tools from model selection to large-scale genomic data offers unprecedented and powerful opportunities to both generate and impartially test evolutionary and mechanistic hypotheses [24] (aligning with an influential recent commentary on ideas in biology [25]). Here, following previous work on mtDNA evolution [19], we adopt this philosophy to explore the mechanisms shaping gene loss across organelles. First, mindful of the dangers of proposing parallels between different organelles [26], we nonetheless hypothesised that the same genetic features would shape retention propensity of genes in mitochondrial and plastid DNA. Such features would predispose a gene to be more or less readily retained in oDNA overall, while the total extent of oDNA retention in a given species is shaped in parallel by functional and metabolic features [23, 27] and evolutionary dynamics (characterised statistically in elegant recent work [28]). We further expect that these genetic features would reflect the above evolutionary tension, between maintaining genetic integrity and retaining the ability to obtain and control machinery, that applies to both organelles [29, 7]. With this general hypothesis in mind, we proceed by taking an impartial, data-driven approach using large-scale genomic data to investigate which features of genes and their protein products predict oDNA gene retention presence (whether any eukaryotes retain a given gene in oDNA) and extent (how commonly an oDNA gene is retained across eukaryotes).

## Results

### Quantifying gene-specific oDNA loss patterns across eukaryotes

To quantitatively explore the features predicting oDNA gene retention, we first define a retention index for a given oDNA gene, measuring its propensity to be retained in oDNA. To this end, we acquired data on organelle gene content across eukaryotes, using 10328 whole mtDNA and 5176 whole ptDNA sequences from NCBI. We curated these data with two different approaches, resembling supervised and unsupervised philosophies, to form consistent records of gene presence/absence by species (see Methods). The supervised approach (manual assignment of ambiguous gene records to a chosen gene label) and the unsupervised approach (all-against-all BLAST comparison of every gene record from the organelle genome database) agreed tightly (Supplementary Fig. S1). Simply counting observations of each gene across species is prone to large sampling bias, as some taxa (notably bilaterians and angiosperms) are much more densely sampled than others. Instead we reconstructed gene loss events using oDNA sequences of modern organisms and an estimated taxonomic relationship between them (see Methods). Motivated by hypercubic transition path sampling [19, 30], we then define the retention index of gene *X* as the number of other genes already lost when gene *X* is lost (results were robust with alternative definition; see below). This retention index, along with the unique patterns of oDNA gene presence/absence and their taxonomic distribution, are illustrated in Fig. 1A (phylogenetic embedding in Supplementary Fig. S2).

**Figure 1:**
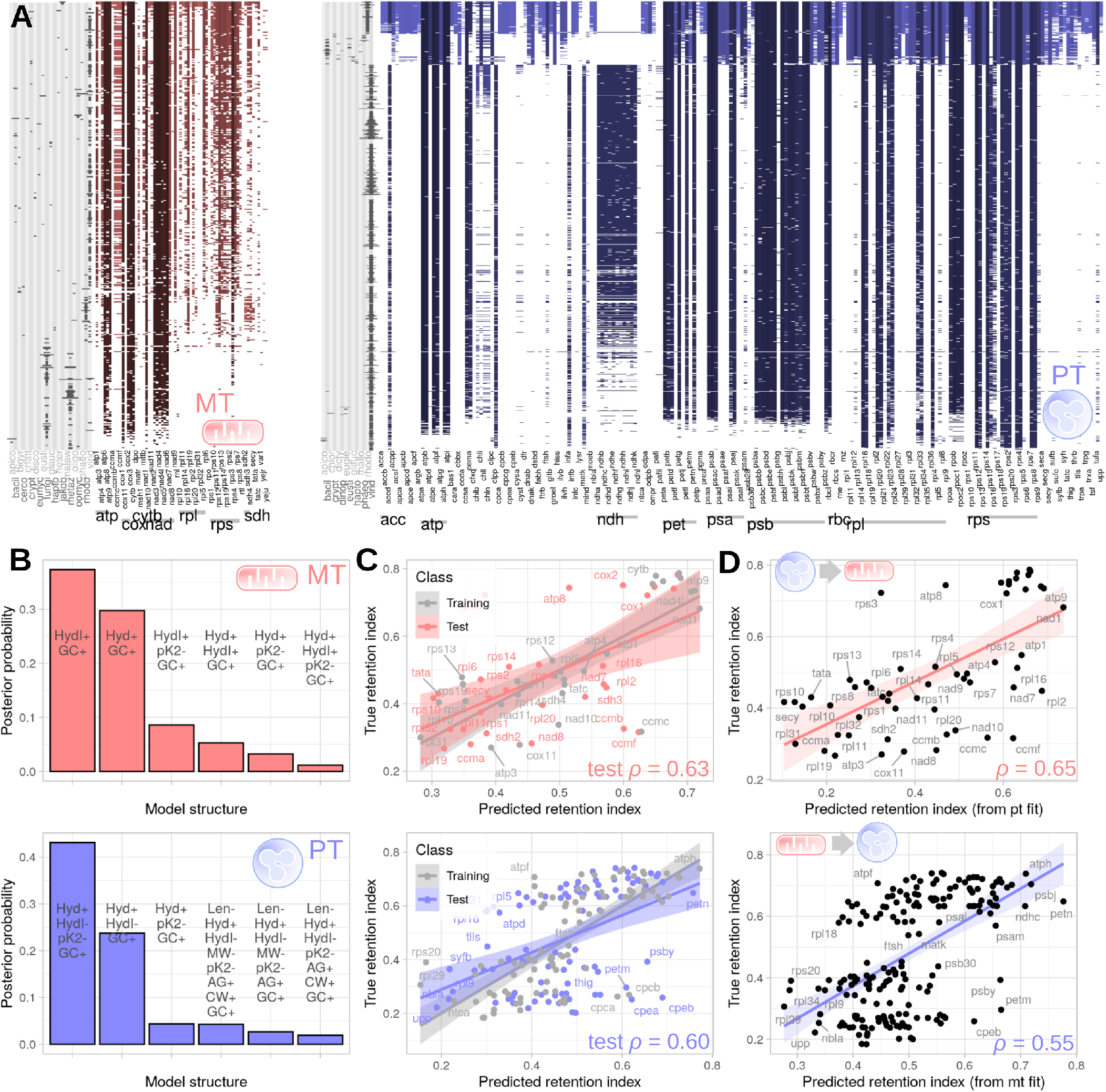
Structure and predictors of oDNA gene retention. (A) Each row of coloured/white pixels is a unique gene presence/absence pattern found in eukaryotic oDNA, where columns are individual oDNA genes. Darker colours correspond to higher values of our assigned retention index for a given gene. Each pattern may be present in many species: grey bars on the left of each row show the number of species with that pattern in a number of eukaryotic clades. The pronounced split in ptDNA patterns reflects the evolutionary pathways represented, for example, by Rhodophyta and Viridiplantae [3]. Sets of genes encoding subunits of notable organelle protein complexes are labelled with grey bars under the horizontal axis. Full set of taxon abbreviations is in Supplementary Text; notable taxa are [metaz]oa, [virid]iplantae, [fungi], [apico]mplexa, [jakob]ida, [rhodo]phyta. (B) Posterior probabilities over the set of features in linear models predicting retention index. Each model structure is given by a set of codes describing its component features. Hydrophobicity (Hyd) or hydrophobicity index (HydI) and GC content (GC) feature in all model structures with the highest posterior probabilities (for priors see Methods). +*/*− give posterior mean signs of associated coefficients in model for retention index. Full feature list: [Hyd]rophobicity, [HydI] hydrophobicity index, [GC] content, [Len]gth, [pK1] carboxyl pKa, [pk2] amino pKa, [MW] molecular weight, [AG/CW] energies of gene expression (Supplementary Text). (C) Prediction of retention index with linear models involving hydrophobicity and GC content. oDNA gene sets are split into training and test sets; trained models predict retention indices well in the independent test sets. (D) Cross-organelle prediction. Linear models trained on mtDNA gene properties predict retention indices of ptDNA genes well, and vice versa.

The retention patterns of genes in mtDNA and ptDNA across eukaryotes show pronounced structure, arguing against a null hypothesis of random gene loss. The several-fold expansion of mtDNA in this study compared to [19] preserves the same structure, with, for example, several *rpl* genes and *sdh[2-4]* commonly lost and *nad[1-6], cox[1-3]* and *cytb* commonly retained. The ptDNA patterns display pronounced clustering, following previous observations [31], with one cluster corresponding broadly to Viridiplantae (typically retaining *ndh* genes) and the other corresponding broadly to brown and red algae, diatoms, and other clades (typically lacking *ndh* genes but retaining more *atp, rps, rpl, psa*, and *psb*). Several ribosomal subunits and *ndhb* are among the most retained in ptDNA, with a second tier involving many *ndh, psa, psb*, and *atp* genes retained in around half our species. Least retained ptDNA genes include other members of *psa, psb, rps*, and *rpl*.

### Cross-organelle symmetry in the prediction of gene retention by hydrophobicity and GC content

We next compiled a set of quantitative properties of genes and their protein products, linked to evolutionary hypotheses about the mechanisms shaping oDNA gene retention [19]. These included gene length and GC content, statistics of encoding and codon usage, and protein hydrophobicity, molecular weight, energy requirements for production, average carboxyl and amino pKa values for amino acid residues, and others (Supplementary Fig. S3). Our quantitative estimates for each feature were averages over a taxonomically diverse sampling of eukaryotic records (see Methods). We used Bayesian model selection to ask which of these properties were most likely to be included in a linear model predicting the retention index of each gene. Following Ref. [19], this approach identifies likely predictors with quantified uncertainty, while acting without prior favouring of any given hypotheses, and automatically guarding against overfitting and the appearance of correlated predictors providing redundant information. In both mtDNA and ptDNA datasets, models where high hydrophobicity and high GC content predict high gene retention were strongly favoured (Fig. 1B). It is well-known that oDNA generally has lower GC content than nuclear DNA, because of the asymmetric mutational pressure arising from the hydrolytic deamination of cytosine to uracil, reducing GC content in the high mutation system of oDNA [32]. However, our results show that higher GC content is relatively favoured between oDNA genes – and so at least partly independently of the general oDNA/nDNA difference [19].

We then tested the capacity of models involving these features to predict the retention index of oDNA genes. We split mtDNA and ptDNA gene sets into 50:50 training and test sets, trained linear models involving hydrophobicity and GC content using the training data, and examined their performance in prediction retention index in the independent test set. Average Spearman correlations were *ρ* = 0.64 and *ρ* = 0.62 for training mt and pt sets respectively, and *ρ* = 0.63 and *ρ* = 0.60 for test mt and pt sets respectively (Fig. 1C). Correlations were higher still (*ρ* > 0.7) when only subunits of core bioenergetic complexes were considered (Supplementary Table S1). Following our hypothesis that the same features predict retention in the two organelle types, we also performed cross-organelle experiments. That is, we trained a hydrophobicity and GC model using mt genes and examined its ability to predict pt gene retention, and vice versa. Strikingly, both organelle gene sets predicted well the other’s retention patterns (*ρ* = 0.65 for pt predicting mt; *ρ* = 0.55 for mt predicting pt; Fig. 1D, Supplementary Table S1). In other words, a simple model trained only using mitochondrial gene data can predict the retention profile of plastid genes, and vice versa.

To relax the assumptions involved in this analysis, including linear modelling, we paralleled this analysis with a range of other regression approaches from data science, including penalised regression and random forests, and using different definitions of retention index (Supplementary Text; Supplementary Fig. S4). We generally observed hydrophobicity and GC content being selected as features with good predictive ability and the capacity to predict one oDNA type’s behaviour from the other, regardless of statistical approach taken (Supplementary Table S1); pKa values were also selected as informative features in some model types (see below).

### Hydrophobicity and protein biochemistry predicts oDNA gene transfer to the nucleus in both organelles

We next asked which properties predict which organelle protein-coding genes are universally transferred to the nucleus across all eukaryotes. To this end, we compiled sets of annotated nDNA and oDNA genes encoding subunits of bioenergetic protein complexes in organelles using a custom pattern matching algorithm and 308 eukaryotic whole genome records from NCBI (see Methods) (Fig. 2A). As expected, GC content in organelle-encoded genes was systematically lower than nuclear-encoded genes. Here, this signal cannot be regarded as a causal mechanism, because it is likely due at least in part to the aforementioned differences in asymmetric mutational pressure between nDNA and oDNA [32, 19]. More interestingly, the hydrophobicity of organelle-encoded genes was systematically higher across taxa (agreeing with recent observations in the mitoribosome [33]), and the carboxyl pKa values of organelle-encoded genes were also systematically higher; other features also differed by encoding compartment (Supplementary Fig. S5). We used Bayesian model selection with a generalised linear model (GLM) using gene properties to predict the encoding compartment (except GC and codon use statistics, due to the possibility of differences therein arising simply due to asymmetric mutation). We found that hydrophobicity and carboxyl pKa consistently appeared in all the model structures with highest posterior probability. Their appearance together in a Bayesian model selection framework suggests that they provide independent information on gene encoding, despite a correlation (albeit rather weak) between the features (Supplementary Fig. S3). GLMs using hydrophobicity and carboxyl pKa, trained using a subset of genes from a given species, were able to to predict the encoding compartment of an independent test set from that species with high performance (True Positive/Negative rates: mt TP 0.90 ±0.17, TN 0.97 ±0.10, pt TP 0.75 ±0.20, TN 0.88 ±0.18, mean and s.d. across species). We also verified that these differences existed within the sets of genes encoding subunits of different organellar complexes (Fig. 2B). We employed a range of classification approaches to quantify these observations, again training on a subset of the observations and testing classification performance on an independent set (Supplementary Fig. S12). Hydrophobicity and pKa values consistently appeared as strong separating terms, with other features including production energy and gene length playing a supporting role (Supplementary Fig. S12). Classification accuracy was typically > 0.8 for all complexes using random forest approaches (Supplementary Table S4).

**Figure 2:**
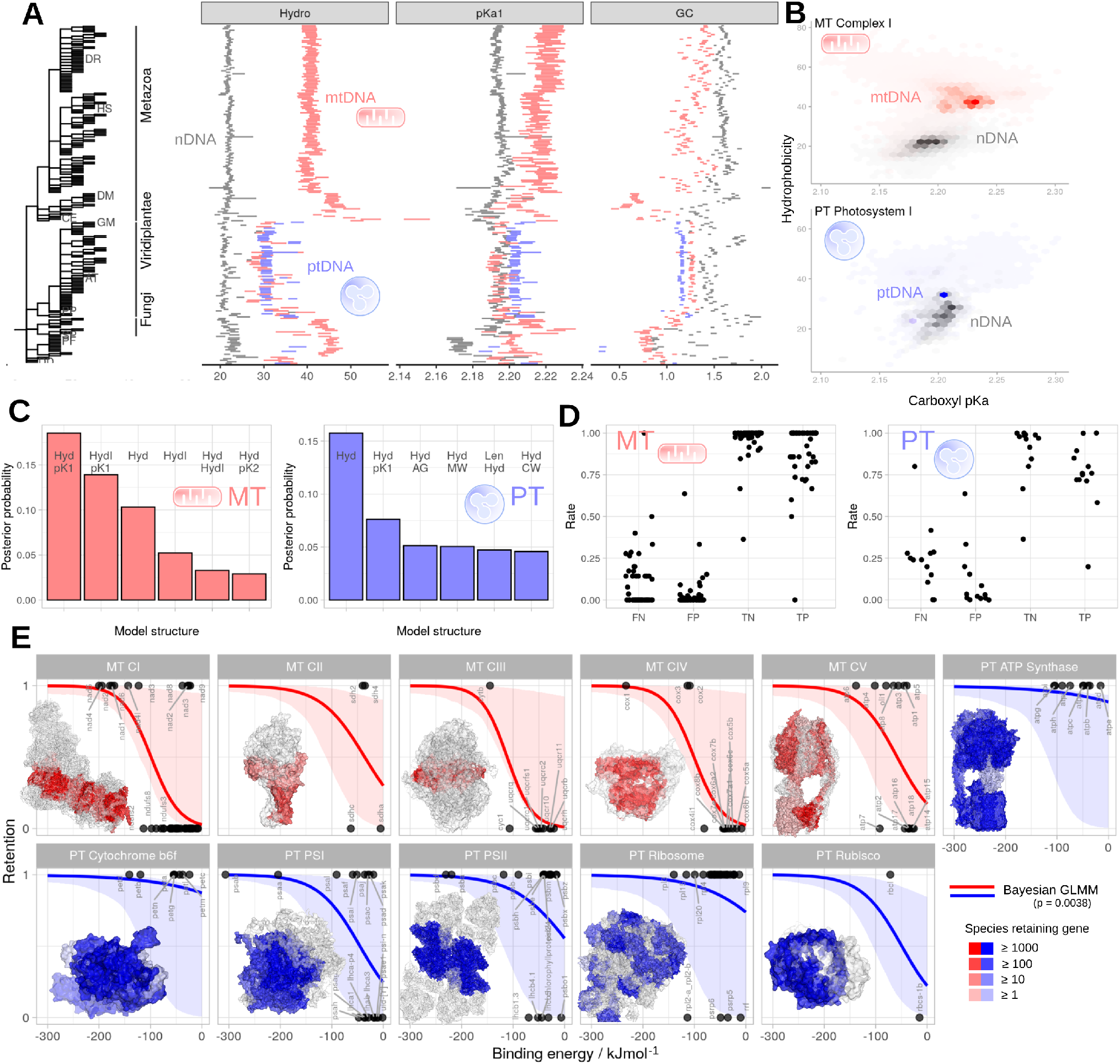
Features predicting encoding compartment. (A) Mean and s.e.m. of selected gene properties for organelle genes encoded in nuclear DNA (grey), mtDNA (red), and ptDNA (blue), in different species (organised by the phylogeny on the left, expanded set in Supplementary Fig. S5). (B) Hydrophobicity and carboxyl pKa of organelle genes encoded in nuclear DNA (red) and oDNA (blue), organised by the protein complex that the gene product occupies (expanded set in Supplementary Fig. S6). (C) Bayesian model selection with a generalised linear model (GLM) framework for features predicting the encoding compartment of a given gene. Posterior probabilities are averaged across independent classifications for individual organisms. Each model structure is given by a set of codes describing its component features; model labels as in Fig. 1. (D) Performance (True/False Positive/Negative) of GLMs involving hydrophobicity and carboxyl pKa on predicting encoding compartment of genes outside the training set. Each set of points corresponds to a model for one organism. (E) Binding energy and encoding compartment. Traces show mean and 95% credible intervals for Bayesian generalised linear mixed model (GLMM) (see Methods for priors). The associated p-value is a frequentist interpretation from bootstrapping, against the null hypothesis of no relationship. Crystal structures are coloured according to the number of species in our dataset that retain the gene for each subunit.

For a subset of organelle-localised gene products, solved crystal structures of their protein complexes allow another property to be quantified: the binding energy statistics of the protein product in its protein complex structure. Previous work qualitatively suggested that genes encoding subunits with high total binding energy (strong binding interactions with neighbouring subunits) and playing central roles in complex assembly pathways were most retained in mtDNA [19, 34, 14]. We used a generalised linear mixed model to quantify and extend this analysis to complexes in both organelle types. We found that total binding energy predicted whether a gene was organelle-encoded in any eukaryotes, with the relationship holding across mitochondria and plastids, though with varying magnitudes in different complexes (Fig. 2C; Supplementary Fig. S7). We verified the absence of pronounced correlation structure between binding energy statistics and hydrophobicity (Supplementary Fig. S8), suggesting that the two features independently contribute to gene retention [19]. Hence, hydrophobicity, amino acid biochemistry, and energetic centrality (linked to colocalisation for redox regulation [14]) predict whether a gene is ever retained in oDNA; of those that are, hydrophobicity and GC content predict the extent of this retention across eukaryotes.

### Independent endosymbiotic genomes show compatible profiles of hydrophobicity and protein biochemistry

Evolutionary history cannot easily be rerun to independently examine these principles. However, the diversity of eukaryotic life provides some existing opportunities to test them. In several eukaryotic species, unicellular endosymbionts that are not directly related to mitochondria or plastids have co-evolved with their ‘host’ species, in many cases involving gene loss and in some cases transfer of genes to the host. Class *Insecta* are known to have several examples of reduced bacterial endosymbionts [35]; other notable examples include the chromatophore, an originally cyanobacterial endosymbiont of *Paulinella* freshwater amoebae [36], the recently discovered *Candidatus Azoamicus ciliaticola*, a denitrifying gammaproteobacterial endosymbiont within a *Plagiopylea* ciliate host [37], and the *Nostoc azollae* symbiont of the *Azolla* water ferns [38].

Not all of these endosymbiotic relationships have been shown to involve gene transfer to the host cell nucleus, although there is evidence for this in the *Paulinella* system [39]. All cases do, however, involve reduction of the endosymbiont genome, as some machinery in the endosymbiont becomes redundant in the symbiotic relationship. In a subset of lost genes, this redundancy arises because host-encoded machinery can fulfil the required function (other genes will be lost without such host-encoded compensation, as their entire function becomes redundant).

For this subset, the same broad principles regarding import of protein machinery may then be expected to hold as in organelles. Such genes are lost as host-encoded machinery removes the need for their local encoding. But such host-encoded machinery must be physically acquired by the endosymbiont, raising similar issues of the mistargeting and import difficulty for hydrophobic gene products as in the organelle case. In tandem, any biochemical pressures influencing the ease of gene expression in the endosymbiont compartment may also be expected to shape retention patterns of this subset of genes. We therefore hypothesised that the principles we find to shape gene retention in mitochondria and plastids would also show a detectable signal in these independent endosymbiotic cases (while expecting a lower magnitude hydrophobicity signal, due to loss of some genes without the requirement for nuclear compensation).

To test this hypothesis, we computed genetic statistics for the genomes of endosymbionts and non-endosymbiotic close relatives (Methods; Supplementary Table S2). The hydrophobicity profile of the endosymbionts in 9 of 10 cases was significantly higher than their non-endosymbiotic relative (Supplementary Text; Fig. 3). Genes retained in the photosynthetic chromatophore also had lower carboxyl pKa values than in a free-living relative; for other endosymbionts, this relationship was reversed, with endosymbiont genes having lower carboxyl pKa values. This is compatible with a possible mechanistic link between the pH of the compartment and the dynamics of gene expression therein (see Discussion).

**Figure 3:**
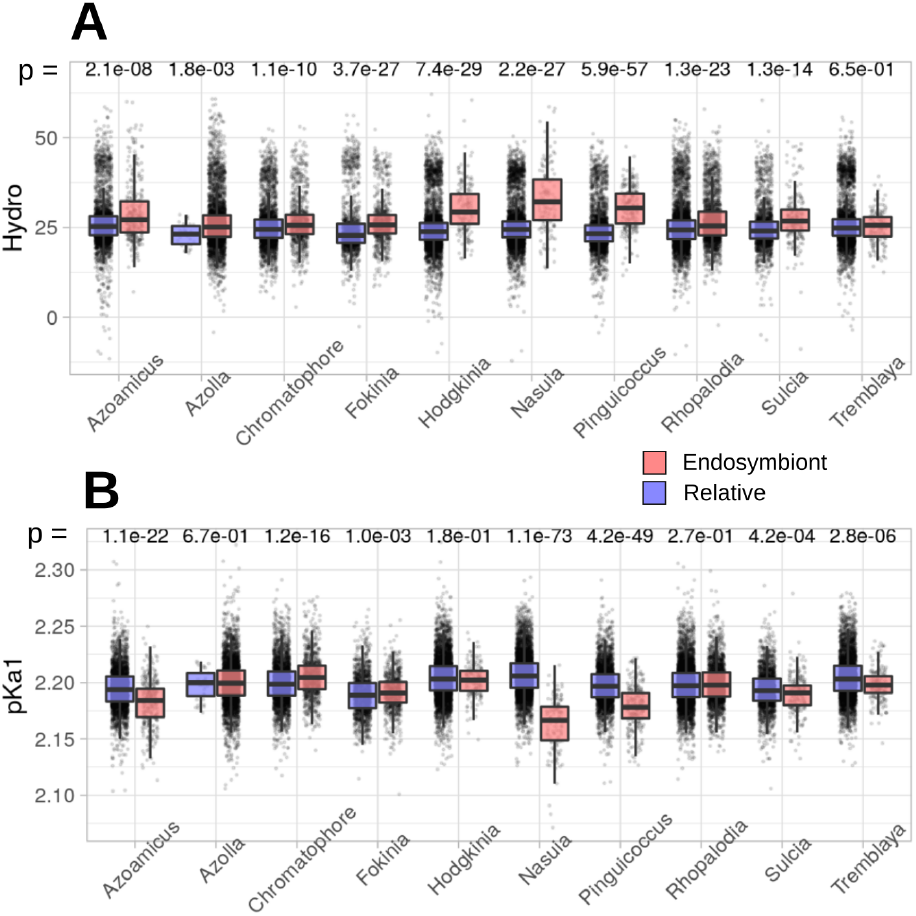
Gene feature profiles in other endosymbionts. Hydrophobicity and carboxyl pKa across genes in endosymbionts (red) and a non-endosymbiotic close relative (blue). p-values are from Wilcoxon rank-sum tests.

Our analysis approach involves several choices of parameter and protocol. To assess the robustness of our findings, we have varied these choices and checked the corresponding change in outputs, described in Supplementary Text and the following figures. The key choices, with figures illustrating their effects, are in gene annotation (supervised or unsupervised; Supplementary Fig. S1), initial selection of features (where we followed existing hypotheses and particularly their summary in [19]) and how to summarise their quantitative values (Supplementary Fig. S9), definition of retention index (Supplementary Table S1; Supplementary Fig. S10), choice of priors in Bayesian model selection (Supplementary Fig. S11), and choice of regression and classification methods: we additionally tested LASSO and ridge regression, and decision trees and random forests for regression and classification (Supplementary Figs. S10 and S12).

## Discussion

To summarise, we have found that hydrophobicity and energetic centrality (the latter linked to colocalisation for redox regulation [14]), with other features of nucleic acid and amino acid biochemistry, predict the prevalence of gene retention to a strikingly symmetric extent in mitochondria, chloroplasts, and independent endosymbionts. It must be underlined that no single mechanism has sole predictive power over this behaviour. As expected in complex biological systems, a combination of factors is likely at play, a situation that has perhaps contributed to the ongoing debate on this topic. Our findings support some previously proposed mechanisms for how selective pressures on gene content may be manifest, while not being incompatible with others (for example, recent theory on the energetic costs of encoding and importing genes [21]). Due to the physical difficulty of importing hydrophobic products or their propensity to be mistargeted to other compartments, hydrophobic gene retention may be favoured [15, 17] (though these mechanisms are not free from debate [18]). The binding energy centrality of a subunit in its protein complex was suggested as a proxy for control over complex assembly, and thus redox processes, aligning with the CoRR (colocalisation for redox regulation) hypothesis [18]. GC content and carboxyl pKa have less established mechanistic hypotheses. The increased chemical stability of GC bonds [40] has been suggested to support the integrity of oDNA in the damaging chemical environment of the organelle. pKa, reflecting the ease of deprotonation of amino acid subgroups for different pH environments, influences the dynamics of peptide formation in translation [41], resulting in pronounced and diverse pH dependence of peptide formation for different amino acids [42]. Speculatively, we thus hypothesise that the synthesis of protein products enriched for higher-pKa amino acids may involve lower kinetic hurdles in the more alkaline pH of mitochondria, plastids, and the chromatophore, favouring the retention of the corresponding genes. The pH within other endosymbionts, which perform less or no proton pumping, is expected to be lower, in which case the opposite pKa trend observed in Fig. 3 follows this pattern. This harnessing of large-scale sequence data with tools from model selection and machine learning has thus generated, and tested, new understanding of the fundamental evolutionary forces shaping bioenergetic organelles, providing quantitative support for several existing hypotheses and suggesting new contributory mechanisms to this important process.

## Materials and Methods

### Source data

We used the mitochondrion and plastid sequences available from NCBI RefSeq [43], and annotated eukaryotic whole genome data also from NCBI. The accessions and references for the endosymbiont/relative pairs are given in Supplementary Table S2. For biochemical and biophysical gene properties, we used the values from [19], described in the Supplementary Text, using BioPython [44] to assign these to given gene sequences. We averaged gene statistics over representative species from a collection of diverse taxa, both using model species (*Homo sapiens, Arabidopsis thaliana, Saccharomyces cerevisiae, Reclinomonas americana, Chondrus crispus, Plasmodium falciparum*) and randomly selected members of different taxa (Supplementary Text; Supplementary Fig. S9). We used crystal structures and associated HTML descriptions from the PDB [45] references 1oco, 1q90, 2h88, 2wsc, 5iu0, 5mdx, 5mlc, 5o31, 5xte, 6cp3, 6fkf. We used PDBePISA [46] to estimate subunit binding energies with two different protocols, both removing ligands and incorporating them into the overall binding energy value for a subunit (Supplementary Text). We used estimated taxonomies from NCBI’s Common Taxonomy Tree tool [47].

### Gene labelling and evolutionary transitions

Gene annotations are inconsistent across such a diverse dataset. For organelle genomes, we used two approaches. In a supervised approach, where the full set of unique labels found was manually curated and assigned a ‘correct’ label based on biological knowledge. In an unsupervised approach, we used BLASTn to perform an all-against-all comparison of all genes in our dataset. We scored each comparison as the proportional length of the region of identity compared to the reference sequence, multiplied by the proportion of identities across that region. Scores over 0.75 were interpreted as ‘hits’ (e.g. 75% identity over the full sequence, or full identity over 75% of the sequence). If more than 25% of appearance of gene label *X* in the BLAST output involved a ‘hit’ with gene labels *Y*, we interpreted *X* and *Y* as referring to the same gene. This process built a set of pairwise identities, which we then resolved interatively into groups of gene labels assumed to refer to the same gene. We then assigned the most prevalent gene label to all members of that group. In each case, we retained only genes that were present in more than ten species in our dataset. For annotated whole genome data, we used pattern matching for gene annotations based on regular expression identifiers to identify nuclear-encoded subunits of organellar protein complexes (expressions in Supplementary Text).

Using these curated gene sets, we assigned ‘barcodes’ of gene presence/absence (binary strings of length *L*, with 0 denoting gene absence and 1 denoting gene presence) to each species in our dataset. Each of these species is a tip on an estimated taxonomic tree describing their putative evolutionary relationship. Assuming that gene loss is rare and gene gain is very rare, we iteratively reconstructed parent barcodes on this tree by assigning a 0 for gene *X* if all descendants lack *X*, and 1 otherwise. We then identified parent-child pairs where the child barcode had fewer genes than the parent (the opposite is impossible by construction). For each such instance, we record the transition from parent barcode to child barcode as a loss event.

### Retention indices

Our simple retention index is defined as follows. Identify the set of transitions that involve the loss of gene *X*. For each transition in this set, count the genes retained by the parent and the genes retained by the child, and take their mean. The retention index is the mean of this quantity over the set of transitions where *X* is lost. The rationale is to characterise the number of genes that have already been lost when *X* is lost. If a transition event involves only the loss of *X*, the parent-child average will report this number minus 1*/*2. If a transition involves the loss of several other genes in parallel with *X*, the average of the before and after counts is used. We also used an alternative retention index without dependence on an assumed evolutionary relationship, described in Supplementary Text.

### Prediction of retention index

We used Bayesian model selection with non-local priors to promote separation of overlapping models [48]; specifically, moment (MOM) priors parameterised so that a signal-to-noise ratio of > 0.2 is anticipated, compatible with previous findings [19]; a beta-binomial(1, 1) prior distribution on the model space, and a minimally informative inverse gamma prior for noise. Further prior information, and the effects of varying them, are given in Supplementary Text and Supplementary Fig. S11.

We implemented the selection process in the R package *mombf*. We additionally used linear modelling penalised using ridge and LASSO protocols, tree-based, and random forest regression, described in the Supplementary Text and implemented using *glmnet, tree*, and *randomForest* packages.

### Classification of subcellular encoding

We used Bayesian model averaging for generalised linear models (GLMs) predicting encoding compartments with priors giving probability 1*/*2 for the inclusion of each parameter, implemented in *BMA*. We then trained GLMs involving hydrophobicity and carboxyl pKa on a training subset of genes for each species. The training subset was the union of a random sample of half the nuclear-encoded genes and half the organelle-encoded genes in each species, with the test set being the complement of this set. We also used decision tree and random forest approaches for the classification task, described in the Supplementary Text. For binding energy values, we used both a Bayesian GLM treating all complexes independently, with t-distributed priors with zero mean, implemented in *arm*; and a Bayesian generalised linear mixed model with flat priors over coefficients, residuals, and covariance structure, implemented in *blme*. These priors were used to overcome convergence issues given the perfect separation of datapoints observed for some protein complexes. Complexes were visualised in PyMOL [49].

### Code and dependencies

Code is written in R, Python, and C, with a wrapper script for bash, and is freely available at github.com/StochasticBiology/odna-loss. The list of libraries used and corresponding citations are in the Supplementary Text.

## Acknowledgments

LR and JMC are supported by the BBSRC via the MIBTP Doctoral Training Scheme. This project has received funding from the European Research Council (ERC) under the European Union’s Horizon 2020 research and innovation programme (Grant agreement No. 805046 (EvoConBiO) to IGJ).

## Supplementary Text

### Materials & Methods

#### Source data

We used the mitochondrion and plastid sequences available from NCBI RefSeq [1], and annotated eukaryotic whole genome data also from NCBI. The accessions and references for the endosymbiont/relative pairs are given in Supplementary Table S2. For biochemical and biophysical gene properties, we used the values from [2], described in the Supplementary Text, using BioPython [3] to assign these to given gene sequences. We averaged gene statistics over representative species from a collection of diverse taxa, both using model species (*Homo sapiens, Arabidopsis thaliana, Saccharomyces cerevisiae, Reclinomonas americana, Chondrus crispus, Plasmodium falciparum*) and randomly selected members of different taxa (Supplementary Text; Supplementary Fig. S9). Codes used in the figures are [Hyd]rophobicity, [HydI] hydrophobicity index, [GC] content, [Len]gth, [pK1] carboxyl pKa, [pK2] amino pKa, [MW] molecular weight, [AG/CW] energies of gene expression. We used crystal structures and associated HTML descriptions from the PDB [4] references 1oco, 1q90, 2h88, 2wsc, 5iu0, 5mdx, 5mlc, 5o31, 5xte, 6cp3, 6fkf. We used PDBePISA [5] to estimate subunit binding energies with two different protocols, both removing ligands and incorporating them into the overall binding energy value for a subunit (Supplementary Text). We used estimated taxonomies from NCBI’s Common Taxonomy Tree tool [6].

#### Prediction of retention index

We used Bayesian model selection with non-local priors to promote separation of overlapping models [7]; specifically, moment (MOM) priors parameterised so that a signal-to-noise ratio of > 0.2 is anticipated, compatible with previous findings [2]; a beta-binomial(1, 1) prior distribution on the model space, and a minimally informative inverse gamma prior for noise. Further prior information, and the effects of varying them, are given in Supplementary Text and Supplementary Fig. S11. We implemented the selection process in the R package *mombf*. We additionally used linear modelling penalised using ridge and LASSO protocols, tree-based, and random forest regression, described in the Supplementary Text and implemented using *glmnet, tree*, and *randomForest* packages.

#### Classification of subcellular encoding

We used Bayesian model averaging for generalised linear models (GLMs) predicting encoding compartments with priors giving probability 1*/*2 for the inclusion of each parameter, implemented in *BMA*. We then trained GLMs involving hydrophobicity and carboxyl pKa on a training subset of genes for each species. The training subset was the union of a random sample of half the nuclear-encoded genes and half the organelle-encoded genes in each species, with the test set being the complement of this set. We also used decision tree and random forest approaches for the classification task, described in the Supplementary Text. For binding energy values, we used both a Bayesian GLM treating all complexes independently, with t-distributed priors with zero mean, implemented in *arm*; and a Bayesian generalised linear mixed model with flat priors over coefficients, residuals, and covariance structure, implemented in *blme*. These priors were used to overcome convergence issues given the perfect separation of datapoints observed for some protein complexes. Complexes were visualised in PyMOL [8].

**Table S1:** Mean linear model regression performance (Spearman’s *ρ* between predicted and observed indices) predicting retention index in test sets for different cases. Non-standard genes (*msh1/muts, matr, mttb*) are removed from mtDNA sets for these experiments. Labels show simple retention index vs barcode retention index; ‘pruned’ dataset (retaining only mt genes from families *nad, sdh, atp, cox, cytb, rp* and pt from *psa, psb, rp, rbc, ndh, atp, pet*) vs unpruned. Each LM uses only GC content and hydrophobicity.

| Method | MT training | MT test | PT training | PT test | PT predicting MT | MT predicting PT |
| --- | --- | --- | --- | --- | --- | --- |
| LM (simple) | 0.64 | 0.63 | 0.62 | 0.60 | 0.65 | 0.55 |
| LM-pruned (simple) | 0.73 | 0.71 | 0.72 | 0.72 | 0.68 | 0.50 |
| LM (barcode) | 0.71 | 0.69 | 0.58 | 0.56 | 0.72 | 0.59 |
| LM-pruned (barcode) | 0.71 | 0.70 | 0.64 | 0.64 | 0.67 | 0.51 |

### Taxon abbreviations

Eukaryotic clades in the mitochondrial dataset in Fig. 1 are [apico]mplexa, [bacill]ariophyta, [bigyr]a, [cerco]zoa, [chatto]nellaceae, [crypto]phyceae, [disco]sea, [eumyc]etozoa, [eusti]gmatophyceae, [fungi], [glauco]cystophyceae, [hapto]phyta, [heter]olobosea, [jakob]ida, [malaw]imonas, [metaz]oa, [oligo]hymenophorea, [oomyc]ota, [phaeo]phyceae, [rhodo]phyta, [virid]iplantae. Clades in the plastid dataset are [apico]mplexa, [bacill]ariophyta, [chlora]rachniophyceae, [crypto]phyceae, [dicty]ochophyceae, [dinop]hyceae, [eugle]nida, [eusti]gmatophyceae, [glauc]ocystophyceae, [hapto]phyta, [mallo]monadaceae, [pelag]omonadales, [phaeo]phyceae, [rhodo]phyta, [virid]iplantae.

### Alternative retention index definitions

In addition to our simple retention index, which relies on an estimated phylogeny linking observations in our dataset, we considered another assumption-free index. Here, we construct the set of unique oDNA presence/absence patterns in our dataset (as in Fig. 1A), and simply count the occurrences *c*_*i*_ of each gene *i* in this dataset. The index is given by log *c*_*i*_ */* max_*j*_ log *c*_*j*_. This index relies on no evolutionary assumptions, and thus cannot account for the evolutionary relationship between sampled species. Considering only the set of unique barcodes goes some way towards accounting for the sampling bias in the dataset (for example, almost all metazoans have the same presence/absence profile, but this profile will only occur once in the unique set). The distribution of this index had substantial structure (as visible in Fig. 1A, and clear, particularly for plastids, in Supplementary Fig. S10), but we do not consider further transformations or more tailored analysis here, instead focusing on the similarity of results with those from the other index.

### Biochemical and biophysical properties of genes and products

Our assignment of biochemical and biophysical properties of genes and their products follows that in Ref. [2]. The **length\*** (in number of amino acids of gene product) and **GC content** (trivially counted) of genes are taken straightforwardly from a sequence. Chemical properties of amino acids were taken from the compilation at http://www.sigmaaldrich.com/life-science/metabolomics/learning-center/amino-acid-reference-chart.html. The **hydrophobicity** and **hydrophobicity index** of a gene product was computed using this compilation (original data from Ref. [9]). **Amine group pK**_*a*_, **carboxyl group pK**_*a*_, and **molecular weight\*** values were calculated using this compilation (original data from [10]).

**Glucose energy costs\*** were computed using the *A*_*glucose*_ metric, based on the absolute nutrient cost required for amino acid biosynthesis, from Ref. [11]. **Craig-Weber energy costs\***, estimating the number of high-energy phosphate bonds and reducing hydrogen atoms required from the cellular energy pool to produce an amino acid, were taken from Ref. [12]. These biochemical properties are summarised in Supplementary Table S5.

**Figure S1:**
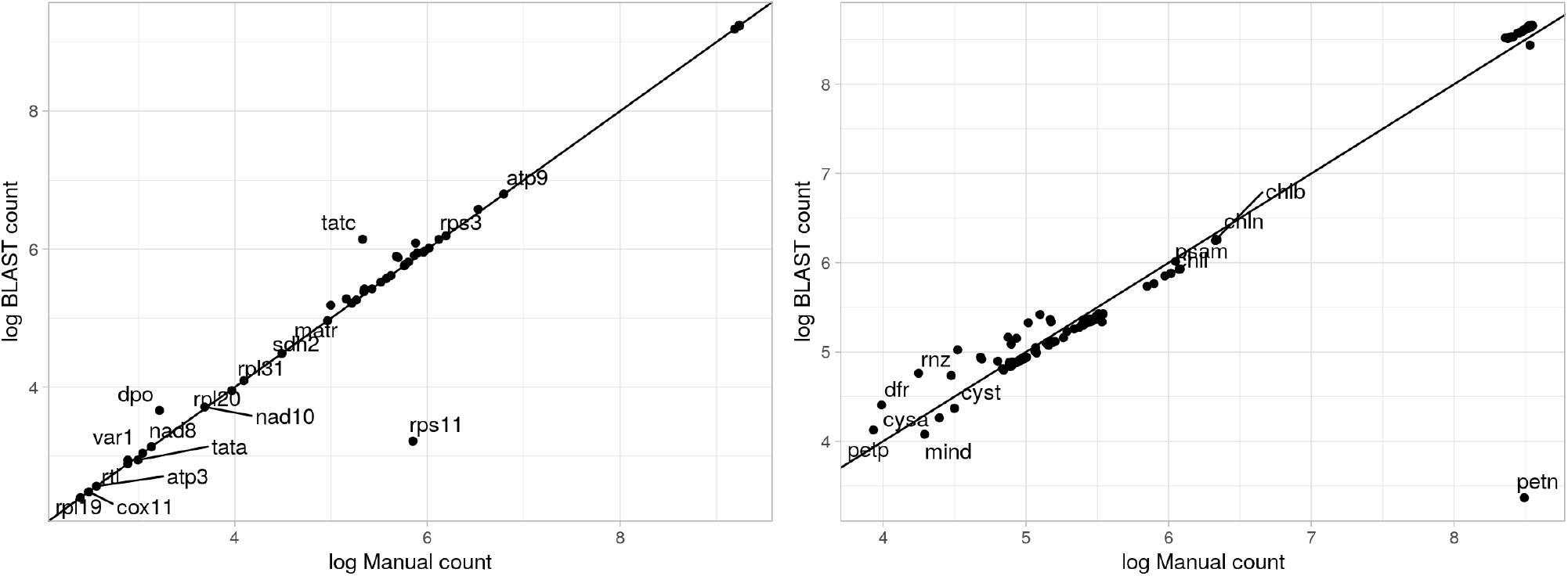
Correlation between gene counts across species derived using manual and BLAST labelling approaches. *r* = 0.9999 for mitochondrial and *r* = 0.9849 for plastid data; discrepancies are largely down to a small number of outliers.

Asterisks denote properties that are *not* averaged over gene length; it was deemed more appropriate to average other properties over genome length to gain a representative measure. To check for artefacts from this interpretation, we performed a (much more computationally demanding) model selection process including both the normalised and un-normalised values for each property; although coverage of individual models was unavoidably low in this procedure, the same consistent observation of GC content and hydrophobicity as important features was observed throughout.

To compute a single value for each statistic of interest, a protocol is required to summarise the many different values seen for a given gene across the species in our dataset. For robustness, we considered several different averaging protocols. First, we averaged gene statistics over a set of model species taken from diverse eukaryotic groups (*Homo sapiens, Arabidopsis thaliana, Saccharomyces cerevisiae, Reclinomonas americana, Chondrus crispus, Plasmodium falciparum*). Second, we randomly selected a member of each clade branching from the eukaryotic group (see clade names above) and averaged over the set containing these random samples. Most statistics were very strongly correlated for these different choices (Fig. S9A). The exception was GC content, which is well known to evolve differently in different clades. To assess the effect of this difference, we ran the model selection process in the text with randomly-sampled averaging protocols. We found that despite differences in GC statistics, the selected models, and the presence of GC within them, remained robust to averaging choice (Fig. S9B).

### Regression for retention index

In addition to the Bayesian linear model approach described in the text, we used a variety of different approaches for retention index regression. These included decision linear modelling with ridge and LASSO penalisation, decision tree regression, and random forest regression. The training, test, and cross-organelle performance of these approaches is given in Table S3.

### Pattern matching for nuclear-encoded organelle genes

We used a combination of positive and negative pattern matching with regular expressions to identify annotations for genes encoding subunits of different organelle complexes. The positive matches required were:

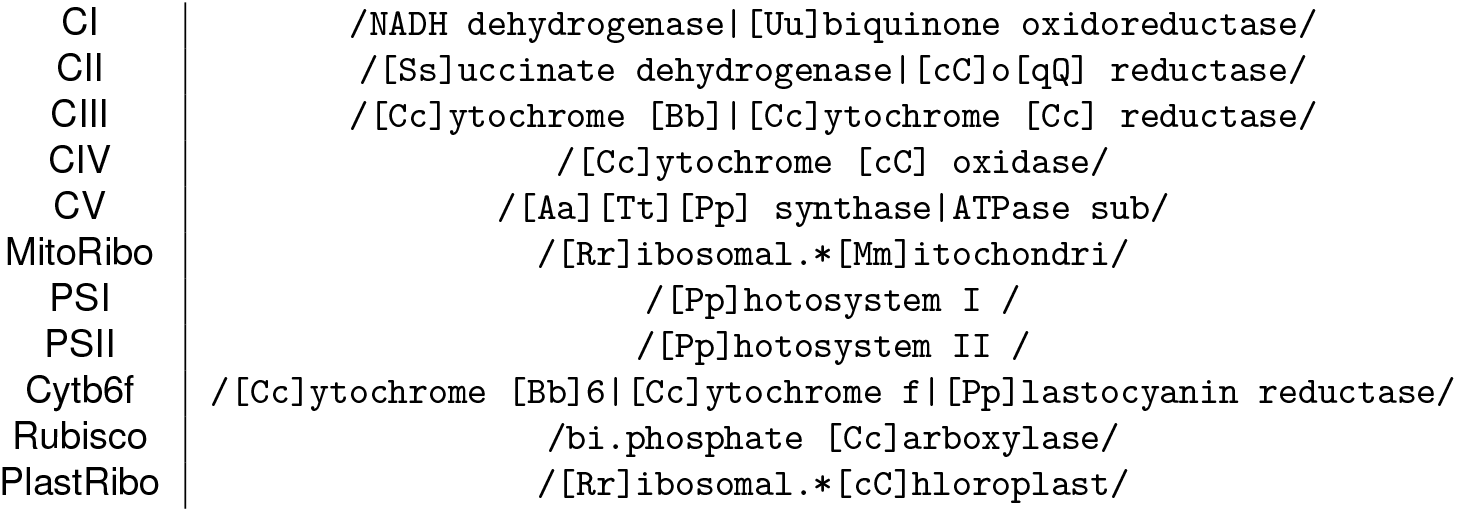

**Figure S2:**
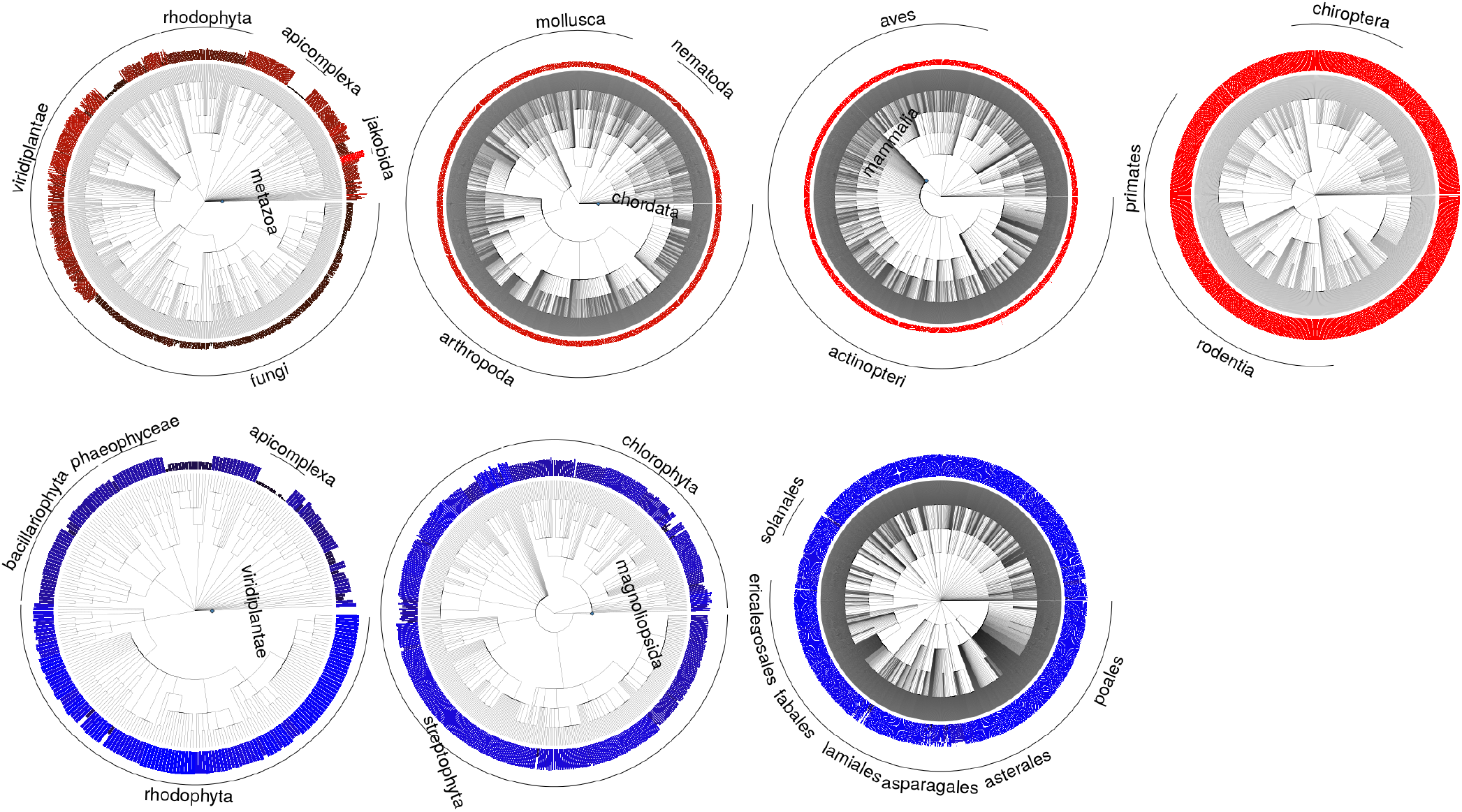
Taxonomic trees for the mt and pt datasets. Blue diamonds give truncation points; associated taxa are expanded in the next rightward tree. Truncated taxa are broadly chosen to reflect those with less diversity in oDNA. Bars illustrate number of retained organelle genes in each species (scale differs in each subtree).

**Figure S3:**
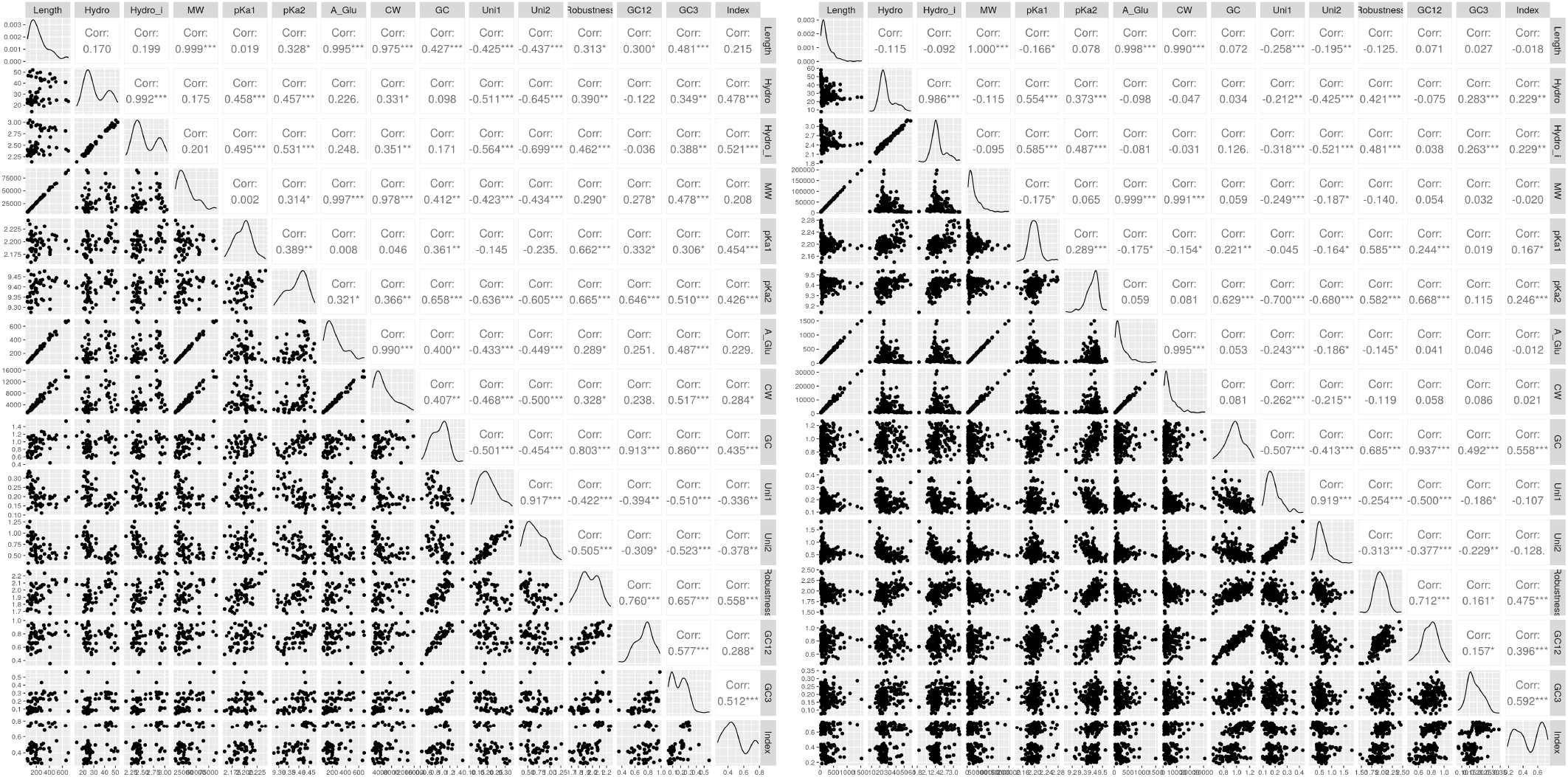
Linear correlations between genetic features and retention index, for mt and pt genes.

**Table S2:** Independent endosymbionts and close free-living relatives. SB, spherical body. ^1^ Relative does invade cells but can survive in oral cavity. ^2^ Partner is not closest sequence found, but is closest annotated sequence in putative phylogeny. ^3^ All closest relatives are intracellular Rickettsiales – relative taken from a sister group. ^4^ Most relatives, including Legionella, are largely intracellular.

| Endosymbiont | NCBI accession | Free-living relative | NCBI accession | References |
| --- | --- | --- | --- | --- |
| *Nasuia deltocephalinicola* | CP013211.1 | *Herbaspirillum seropedicae* | CP002039.1 | [14] |
| *Ca. Sulcia muelleri* | CP001981.1 | *Porphyromonas gingivalis*<sup>1</sup> | AE015924.1 | [15] |
| *Ca. Tremblaya phenacola* | CP003982.1 | *Sodalis praecaptivus* | CP006569.1 | [16] |
| *Rhopalodia gibberula* SB | AP018341.1 | *Cyanothece* sp. *PCC 8801* | CP001287.1 | [17] |
| *Ca. Hodgkinia cicadicola* | CP008699 | *Rhizobium etli* | CP007641.1 | [18] |
| *Ca. Pinguicoccus supinus* | CP039370.1 | *Coralimargarita akajimensis*<sup>2</sup> | CP001998.1 | [19] |
| *Ca. Fokinia solitaria* | CP025989.1 | *Pelagibacter ubique*<sup>3</sup> | CP000084.1 | [20] |
| *Paulinella chromatophore* | CP000815.1 | *Synechococcus* *PCC 7002* | CP000951 | [21] |
| *Ca. Azoamicus ciliaticola* | NZ_LR794158.1 | *Legionella clemsonensis*<sup>4</sup> | NZ_CP016397 | [22] |
| *Nostoc azollae* | CP002059.1 | *Raphidiopsis brookii* | ACYB01000001.1 | [23] |

With the following patterns (split for formatting) required to be absent:

~~~
/assembly|alternative|containing|dependent|chaperone|kinase|NADH-cytochrome|coupling|maturase/
/vacuolar|biogenesis|repair|LOW QUALITY PROTEIN|synthetase|activator|reticulum|activase/
/synthesis|lyase|like| non|transporting|lipid|autoinhibited|membrane|type|required/
/QUALITY|precursor|inhibitor|proteasomal|proteasome|E1|various|regulatory|Clp/
/calcium|vesicle|b-245|b5|WRNIP|AAA|Cation|family|remodelling/
~~~

The outputs of this approach were manually verified to include genes encoding subunits physically present in their corresponding complex, while excluding assembly factors, regulatory factors, synthesis factors, unrelated enzymes, and other false positives.

### Classification for compartment

We also considered decision tree and random forest approaches for the organelle/nuclear encoding compartment classification problem; performance is shown in Table S4, with illustrations in Fig. S12.

### Binding energy calculations

We used PDBePISA [5] to calculate interaction energies between different protein subunits and ligands in crystal structures. We summed the interaction energies over all interfaces between a given subunit and its partners to compute a total energetic centrality statistic for each subunit. Several choices of representation are possible for these calculations. Ligands can be ignored, so that only interaction energies of interfaces directly linking protein subunits are considered. Alternatively, bonds to ligands can be included as contributing to a given subunit’s total binding energy. We primarily considered the mean energy per interface, including ligands, for each subunit, but also verified that our detected relationship existed for different choices including total energy over interfaces.

### Endosymbionts and relatives

We considered a range of endosymbionts highlighted in a comprehensive recent review [13]. For each we sought to identify a close free-living relative. In some cases all closest relatives of an endosymbiont themselves adopted a largely or obligate intracellular lifestyle; in these cases we tried to identify the closest relative that was at least capable of free-living (Table S2).

### Packages and libraries

Our pipeline uses the following R packages: ape [24], arm [25], blme [26], BMA [27], caper [28], cowplot [29], e1071 [30], geiger [31], GGally [32], ggnewscale [33], ggplot2 [34], ggpubr [35], ggpval [36], ggrepel [37], ggtree [38], ggtreeExtra [39], glmnet [40], gridExtra [41], hexbin [42], igraph [43], lme4 [44], logistf [45], mombf [46], nlme [47], phangorn [48], phytools [49], randomForest [50], stringdist [51], stringr [52], and tree [53].

We also use BioPython [3] for parsing sequences and computing gene statistics, PyMOL [8] for visualisation, and BLAST [54] for sequence comparisons.

**Table S3:** Mean regression performance (Spearman’s *ρ* between predicted and observed indices) predicting retention index with different approaches. Non-standard genes (*msh1/muts, matr, mttb*) are not removed for these experiments. Tree, decision tree regression; LM, linear model; Ridge, ridge regression; LASSO, LASSO regression; RF, random forest regression. All genetic features included by default; ‘reduced’ corresponds to models involving only GC content and hydrophobicity. ‘Cross’ refers to cross-organelle experiments where mt training is used to predict pt test and vice versa (N/A, not applicable: no test set within training organelle).

| Method | MT training | MT test | PT training | PT test | PT predicting MT | MT predicting PT |
| --- | --- | --- | --- | --- | --- | --- |
| Tree | 0.79 | 0.40 | 0.82 | 0.45 | 0.54 | 0.33 |
| LM | 0.70 | 0.43 | 0.71 | 0.66 | 0.52 | 0.25 |
| Tree-reduced | 0.73 | 0.48 | 0.75 | 0.45 | 0.55 | 0.39 |
| LM-Reduced | 0.58 | 0.52 | 0.61 | 0.61 | 0.54 | 0.48 |
| Ridge | 0.68 | 0.39 | 0.66 | 0.71 | 0.57 | 0.41 |
| LASSO | 0.63 | 0.44 | 0.66 | 0.71 | 0.57 | 0.37 |
| SVR | 0.81 | 0.46 | 0.77 | 0.62 | 0.62 | 0.34 |
| RF | 0.92 | 0.48 | 0.95 | 0.62 | 0.62 | 0.45 |
| RF-Reduced | 0.88 | 0.50 | 0.92 | 0.51 | 0.57 | 0.50 |
| RF-Cross | 0.94 | N/A | 0.96 | N/A | 0.62 | 0.56 |
| RF-Cross-Reduced | 0.90 | N/A | 0.92 | N/A | 0.55 | 0.59 |

**Table S4:** Nuclear-organelle classification performance (proportion of test set assigned to correct compartment), by organelle complex, with different approaches (tree, decision tree; RF, random forest). Complexes are labelled with regular expressions describing their gene labels. All genetic features included by default; ‘reduced’ corresponds to models involving only GC content and hydrophobicity. ‘Cross’ refers to cross-organelle experiments where mt training is used to predict pt test and vice versa. Balance gives the proportion of genes encoded in one compartment (may fluctuate slightly due to different subsamples being used in model construction): N/A, not applied to cross-organelle classification.

| Complex | Model type | Training | Test | Balance | Complex | Model type | Training | Test | Balance |
| --- | --- | --- | --- | --- | --- | --- | --- | --- | --- |
| nad[0-9] | tree | 0.99 | 0.99 | 0.10 | nad[0-9] | RF | 1.00 | 1.00 | 0.10 |
| sdh[0-9] | tree | 0.97 | 0.91 | 0.66 | sdh[0-9] | RF | 1.00 | 0.95 | 0.68 |
| cytb | tree | 0.99 | 0.99 | 0.18 | cytb | RF | 1.00 | 0.99 | 0.18 |
| cox[0-9] | tree | 1.00 | 0.99 | 0.09 | cox[0-9] | RF | 1.00 | 0.99 | 0.09 |
| atp[0-9] | tree | 0.98 | 0.96 | 0.16 | atp[0-9] | RF | 1.00 | 0.98 | 0.16 |
| (MT) rp[sl] | tree | 0.88 | 0.85 | 0.69 | (MT) rp[sl] | RF | 1.00 | 0.92 | 0.69 |
| psa[a-x] | tree | 0.99 | 0.99 | 0.03 | psa[a-x] | RF | 1.00 | 0.99 | 0.03 |
| psb[a-z] | tree | 1.00 | 0.99 | 0.01 | psb[a-z] | RF | 1.00 | 1.00 | 0.01 |
| atp[a-z] | tree | 0.98 | 0.97 | 0.12 | atp[a-z] | RF | 1.00 | 0.99 | 0.12 |
| pet[a-z] | tree | 1.00 | 0.99 | 0.01 | pet[a-z] | RF | 1.00 | 0.99 | 0.01 |
| rbc | tree | 0.99 | 0.97 | 0.07 | rbc | RF | 1.00 | 0.98 | 0.07 |
| (PT) rp[sl] | tree | 0.99 | 0.99 | 0.02 | (PT) rp[sl] | RF | 1.00 | 0.99 | 0.02 |
| nad[0-9] | tree-reduced | 0.99 | 0.99 | 0.10 | nad[0-9] | RF-reduced | 1.00 | 0.99 | 0.10 |
| sdh[0-9] | tree-reduced | 0.97 | 0.92 | 0.66 | sdh[0-9] | RF-reduced | 1.00 | 0.93 | 0.66 |
| cytb | tree-reduced | 0.98 | 0.97 | 0.18 | cytb | RF-reduced | 1.00 | 0.98 | 0.19 |
| cox[0-9] | tree-reduced | 0.98 | 0.98 | 0.09 | cox[0-9] | RF-reduced | 1.00 | 0.98 | 0.09 |
| atp[0-9] | tree-reduced | 0.92 | 0.91 | 0.16 | atp[0-9] | RF-reduced | 1.00 | 0.92 | 0.16 |
| (MT) rp[sl] | tree-reduced | 0.79 | 0.76 | 0.69 | (MT) rp[sl] | RF-reduced | 1.00 | 0.77 | 0.69 |
| psa[a-x] | tree-reduced | 0.98 | 0.97 | 0.03 | psa[a-x] | RF-reduced | 1.00 | 0.97 | 0.03 |
| psb[a-z] | tree-reduced | 0.99 | 0.99 | 0.01 | psb[a-z] | RF-reduced | 1.00 | 0.99 | 0.01 |
| atp[a-z] | tree-reduced | 0.91 | 0.90 | 0.12 | atp[a-z] | RF-reduced | 1.00 | 0.91 | 0.12 |
| pet[a-z] | tree-reduced | 0.99 | 0.99 | 0.01 | pet[a-z] | RF-reduced | 1.00 | 0.99 | 0.01 |
| rbc | tree-reduced | 0.96 | 0.93 | 0.06 | rbc | RF-reduced | 1.00 | 0.94 | 0.07 |
| (PT) rp[sl] | tree-reduced | 0.98 | 0.98 | 0.02 | (PT) rp[sl] | RF-reduced | 1.00 | 0.98 | 0.02 |
| All PT | tree-cross | 0.94 | 0.80 | N/A | All PT | RF-cross | 1.00 | 0.60 | N/A |
| All MT | tree-cross | 0.98 | 0.82 | N/A | All MT | RF-cross | 1.00 | 0.79 | N/A |
| All PT | tree-cross-reduced | 0.94 | 0.56 | N/A | All PT | RF-cross-reduced | 1.00 | 0.47 | N/A |
| All MT | tree-cross-reduced | 0.97 | 0.81 | N/A | All MT | RF-cross-reduced | 1.00 | 0.82 | N/A |

**Figure S4:**
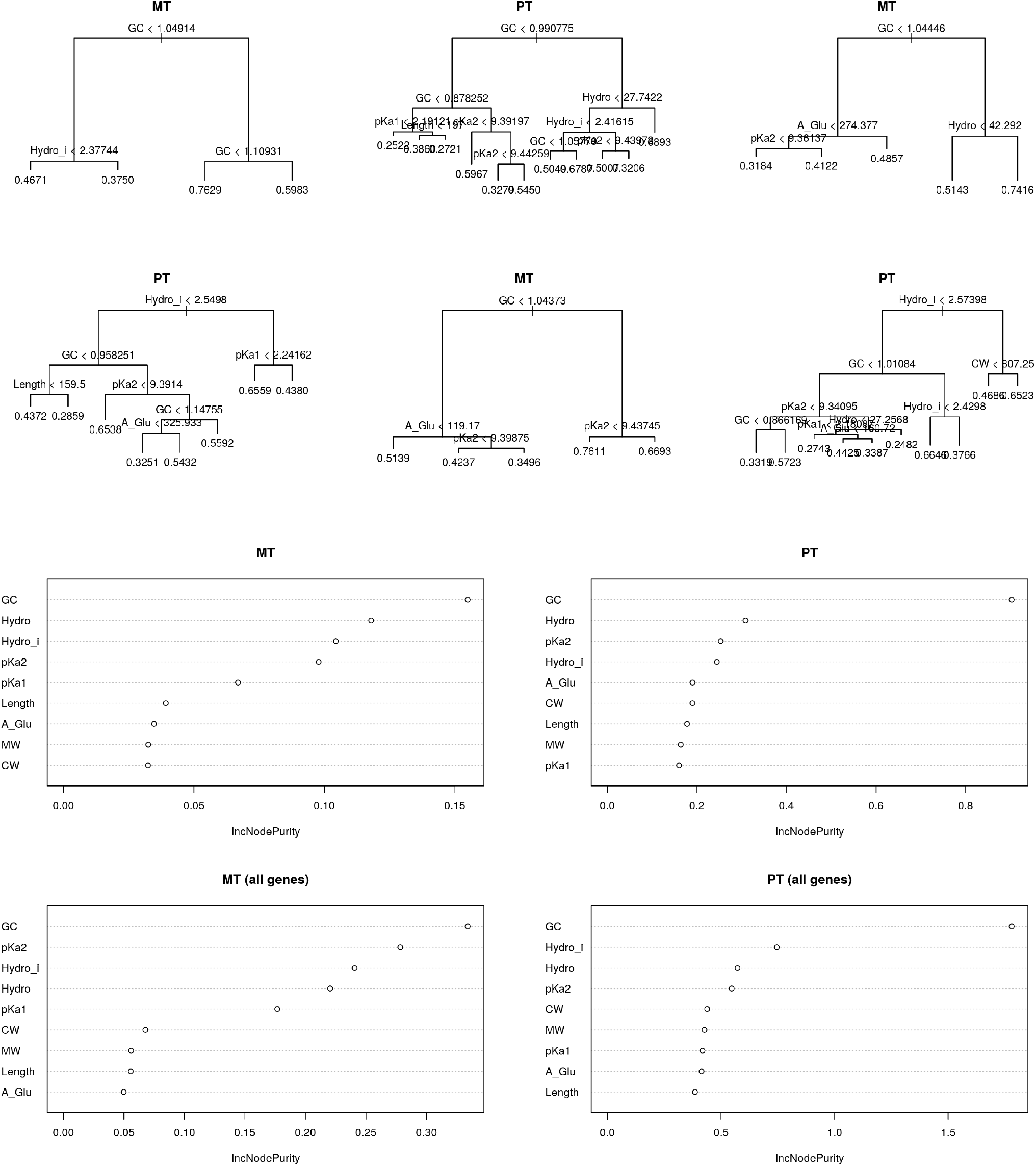
Decision tree and random forest regression for retention index. (top) a set of trees learned to predict retention for different training-test splits, showing the dominant role of GC content and hydrophobicity as predictive features. (bottom) variance improvement plots for random forest regression of the same task, illustrating the importance of each feature in the predictive outcome.

**Figure S5:**
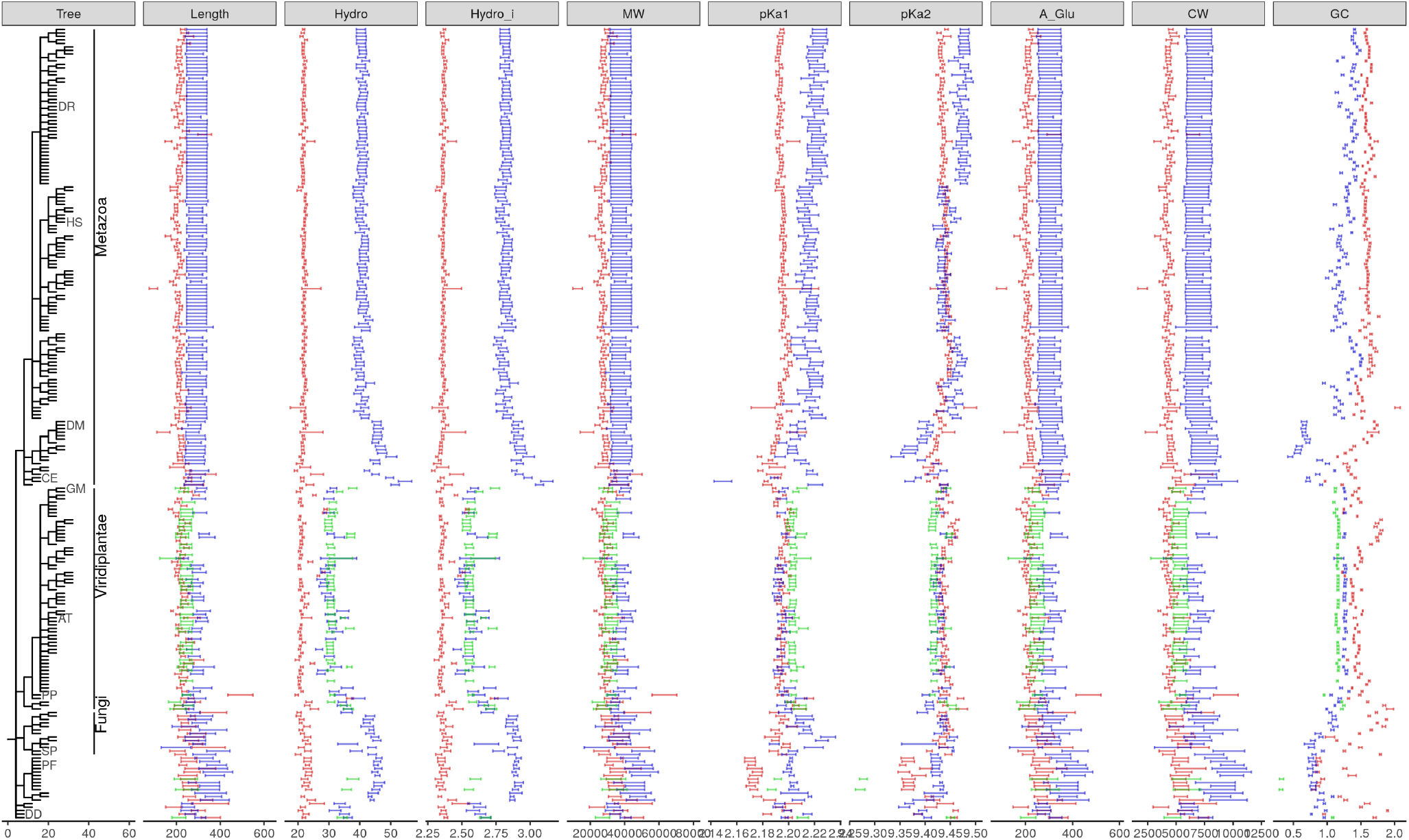
Statistics of genes encoded in the nucleus (red), mitochondrion (blue), or plastid (green) compartments. Bars give mean and s.e.m. for each species; phylogeny shows the relationship between species. Specific model species labelled by initials: *Danio rerio, Homo sapiens, Drosophila melanogaster, Caenorhabditis elegans, Glycine max, Arabidopsis thaliana, Physcomitrella patens, Schizosaccharomyces pombe, Plasmodium falciparum, Dictyostelium discoideum*.

**Table S5:** Amino acid properties used in model selection. Numerical values of the properties described in the text. Qauantities are unitless unless specific. See text for sources.

|  |  | Hydro | Hydro.I | Mol weight / Da | pKa1 | pKa2 | Aglucose | CWEnergy |
| --- | --- | --- | --- | --- | --- | --- | --- | --- |
| Ala | A | 41 | 3 | 89.1 | 2.34 | 9.69 | 0.5 | 12.5 |
| Arg | R | -14 | 1 | 174.2 | 2.17 | 9.04 | 1.39 | 18.5 |
| Asn | N | -28 | 1 | 132.12 | 2.02 | 8.8 | 0.79 | 4 |
| Asp | D | -55 | 1 | 133.11 | 1.88 | 9.6 | 0.61 | 1 |
| Cys | C | 49 | 3 | 121.16 | 1.96 | 10.28 | 0.75 | 24.5 |
| Gln | Q | -10 | 2 | 146.15 | 2.17 | 9.13 | 0.92 | 9.5 |
| Glu | E | -31 | 1 | 147.13 | 2.19 | 9.67 | 0.86 | 8.5 |
| Gly | G | 0 | 2 | 75.07 | 2.34 | 9.6 | 0.31 | 14.5 |
| His | H | 8 | 2 | 155.16 | 1.82 | 9.17 | 1.46 | 33 |
| Ile | I | 99 | 4 | 131.18 | 2.36 | 9.6 | 1.21 | 20 |
| Leu | L | 97 | 4 | 131.18 | 2.36 | 9.6 | 1.21 | 33 |
| Lys | K | -23 | 1 | 146.19 | 2.18 | 8.95 | 1.31 | 18.5 |
| Met | M | 74 | 4 | 149.21 | 2.28 | 9.21 | 1.25 | 18.5 |
| Phe | F | 100 | 4 | 165.19 | 1.83 | 9.13 | 1.84 | 63 |
| Pro | P | -46 | 1 | 115.13 | 1.99 | 10.6 | 0.99 | 12.5 |
| Ser | S | -5 | 2 | 105.09 | 2.21 | 9.15 | 0.49 | 15 |
| Stop | X | - | - | - | - | - | - | - |
| Thr | T | 13 | 2 | 119.12 | 2.09 | 9.1 | 0.69 | 6 |
| Trp | W | 97 | 4 | 204.23 | 2.83 | 9.39 | 2.39 | 78.5 |
| Tyr | Y | 63 | 3 | 181.19 | 2.2 | 9.11 | 1.77 | 56.5 |
| Val | V | 76 | 4 | 117.15 | 2.32 | 9.62 | 0.96 | 25 |

**Figure S6:**
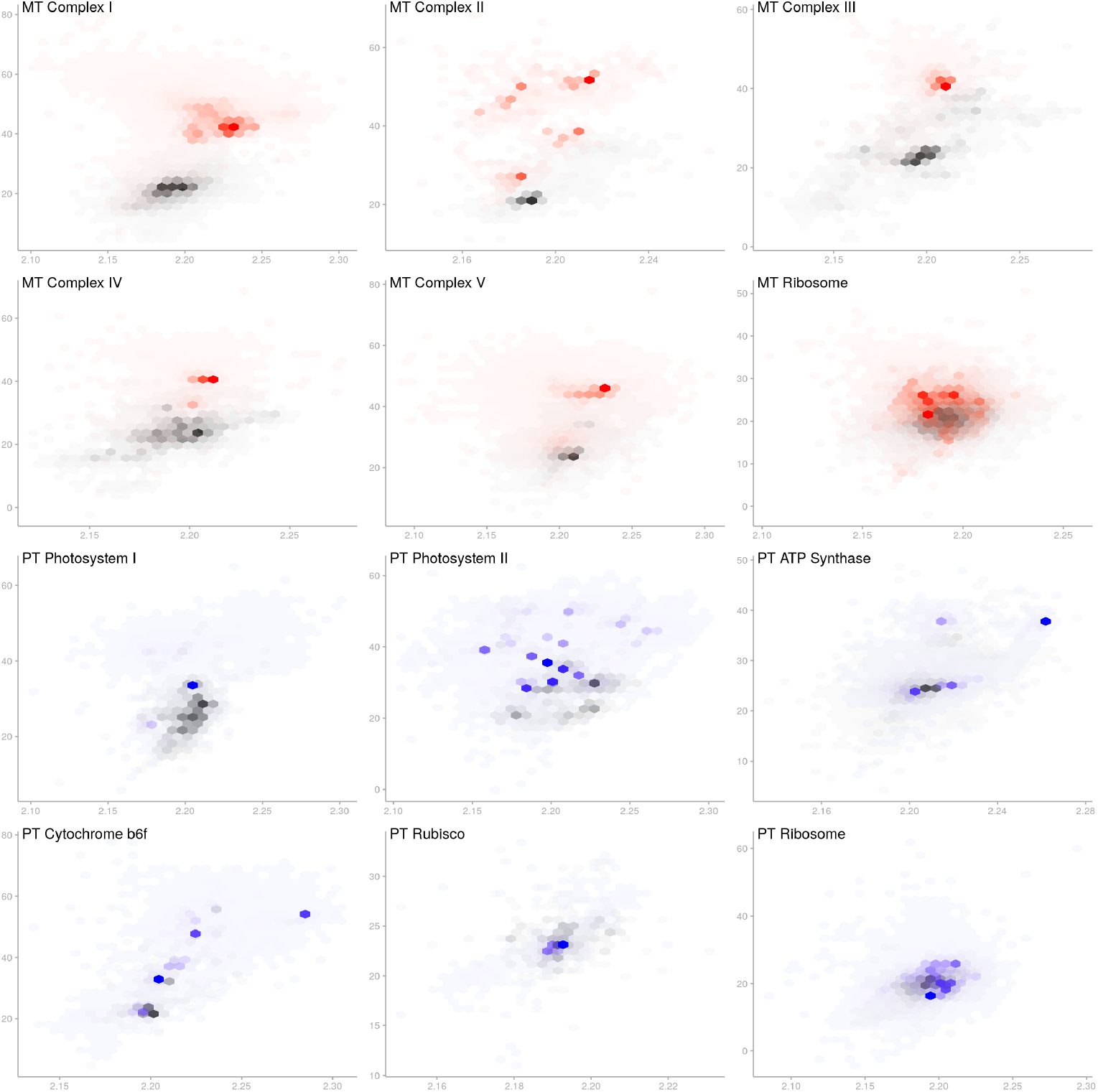
Hydrophobicity and carboxyl pKa for nuclear- and organelle-encoded complex subunits.

**Figure S7:**
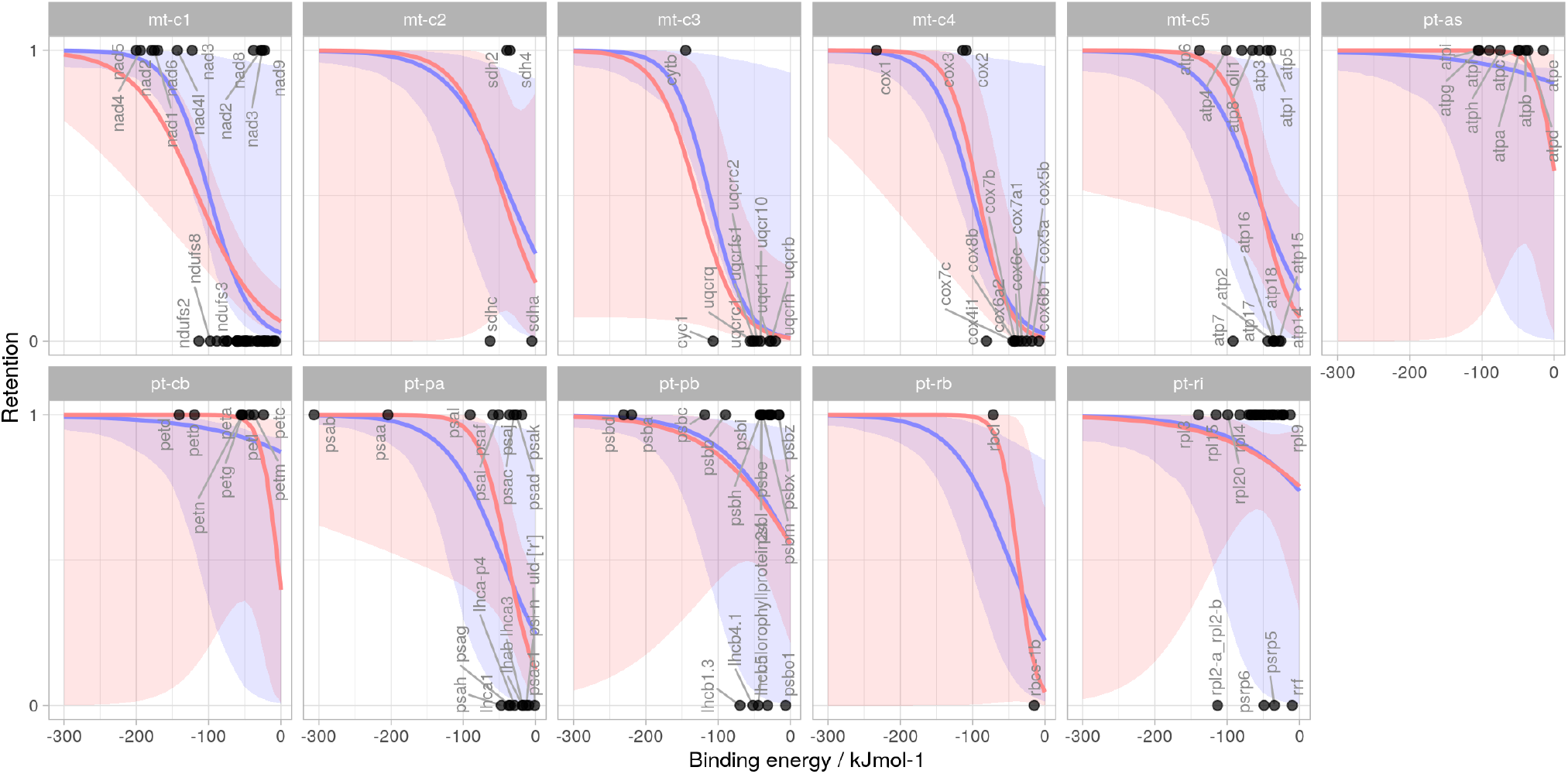
Comparison of Bayesian generalised linear model (GLM) and generalised linear mixed model (GLMM) for binding energy-retention relationship. The GLM approach (red) treats each complex independently; the GLMM (blue) describes complex-specific changes to an overall trend. Frequentist p-values against the null hypothesis of no relationship are 0.00047 (GLM) and 0.0038 (GLMM).

**Figure S8:**
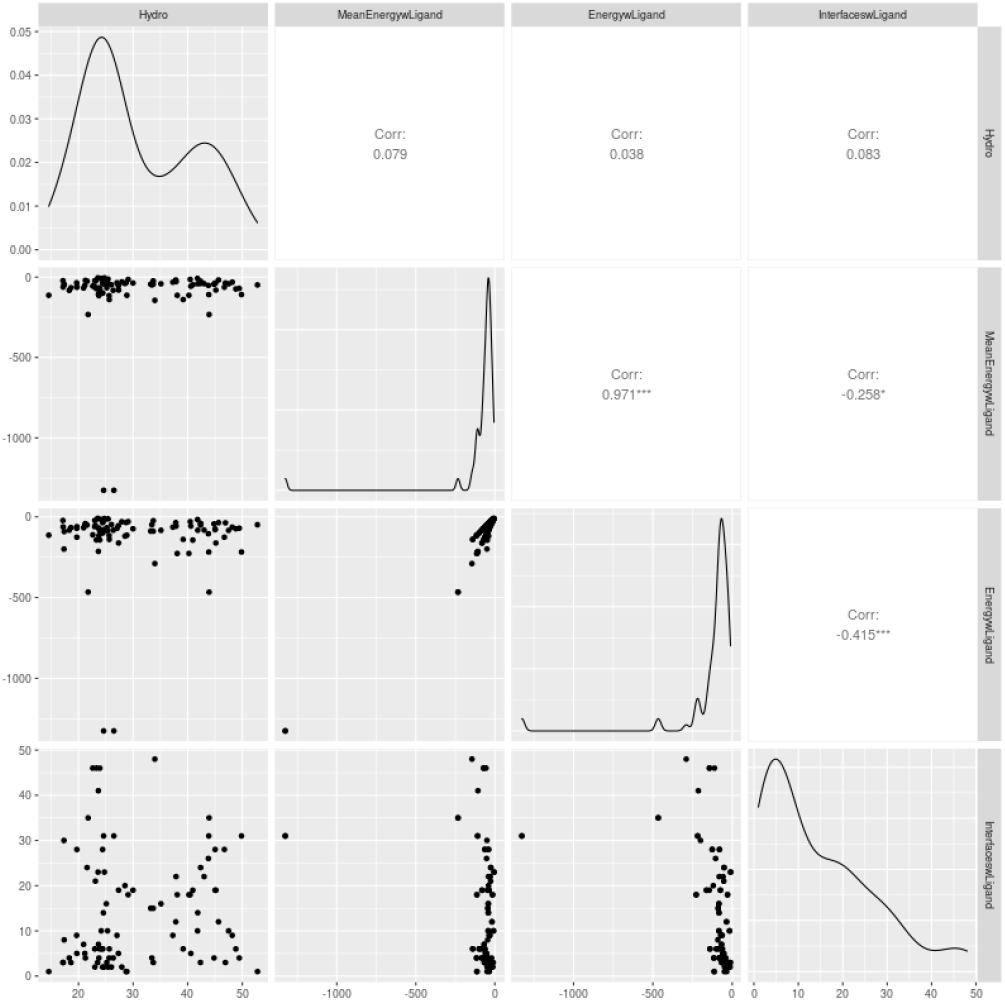
Little correlation between hydrophobicity and energetic centrality across gene products involved in the complexes studied.

---

**Figure S9:**
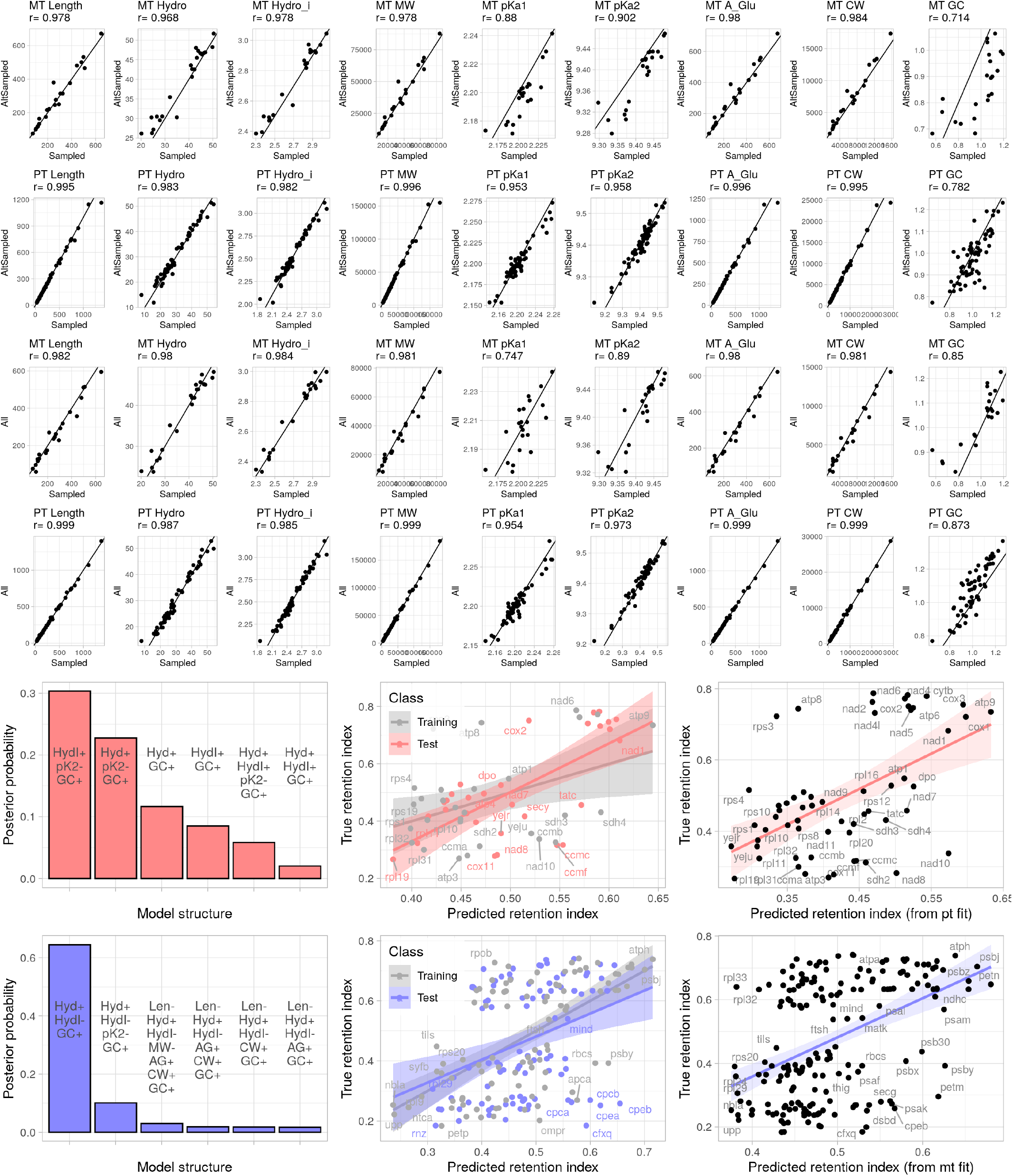
Effect of different averaging protocols to summarise gene statistics. (top) Correlations between our model systems average and (A) average across randomly sampled species from different clades and (B) average across all species in the dataset (expected to be highly weighted towards bilaterians and angiosperms). (bottom) Result of model selection and testing process with the random species averaging protocol.

**Figure S10:**
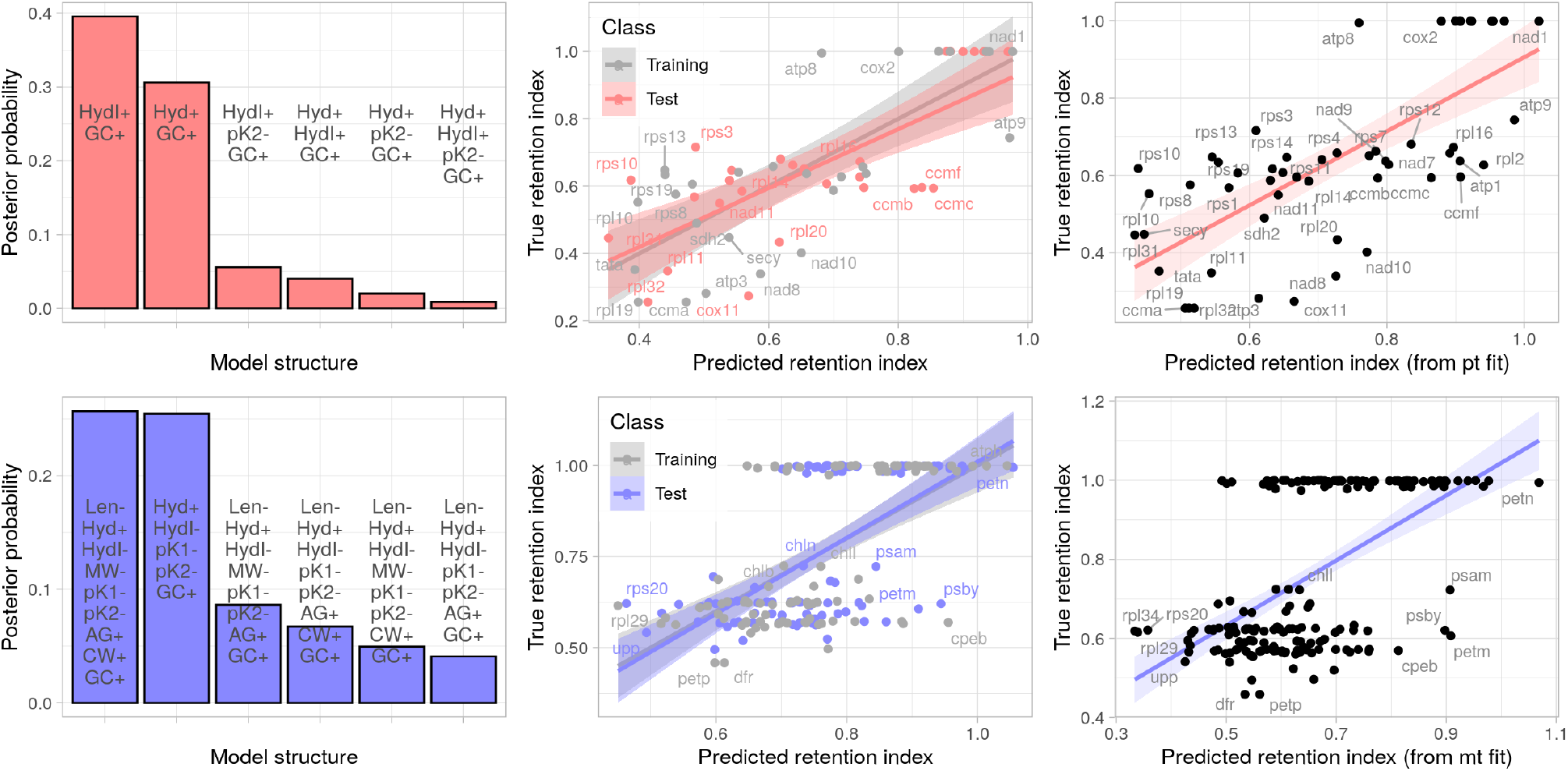
Model selection and regression for the barcode-based retention index reflects the outcomes from the simple retention index (Fig. 1); see also Table S1.

**Figure S11:**
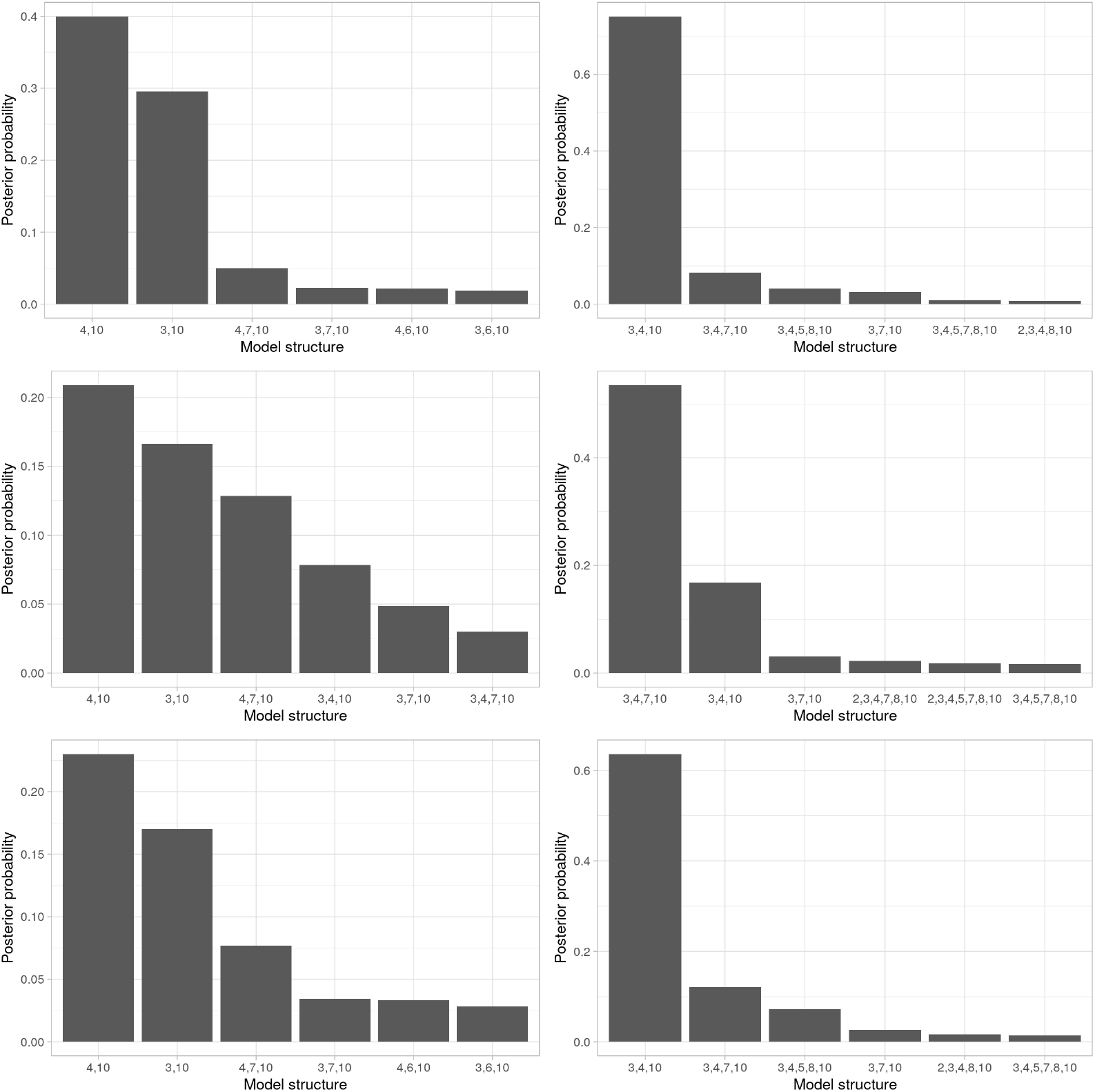
Bayesian model selection for linear models predicting retention index, with different priors from the default choices in the main text. Left, mitochondrial; right, plastid data. Top, inverse moment (iMOM) prior with *τ* = 0.133 and beta-binomial(1,1) prior over models. Centre, moment (MOM) prior with *τ* = 0.348 and uniform prior over models. Bottom, IMOM prior with *τ* = 0.133 and uniform prior over models. MOM vs iMOM changes structure of non-local priors; model priors assign different prior weights to overall model structures. Features appearing in models are: 1 (intercept); 2 (length); 3 (hydrophobicity); 4 (hydrophobicity index); 5 (molecular weight); 6 (amino pKa); 7 (carboxyl pKa); 8 (glucose assembly energy); 9 (alternate assembly energy); 10 (GC content).

**Figure S12:**
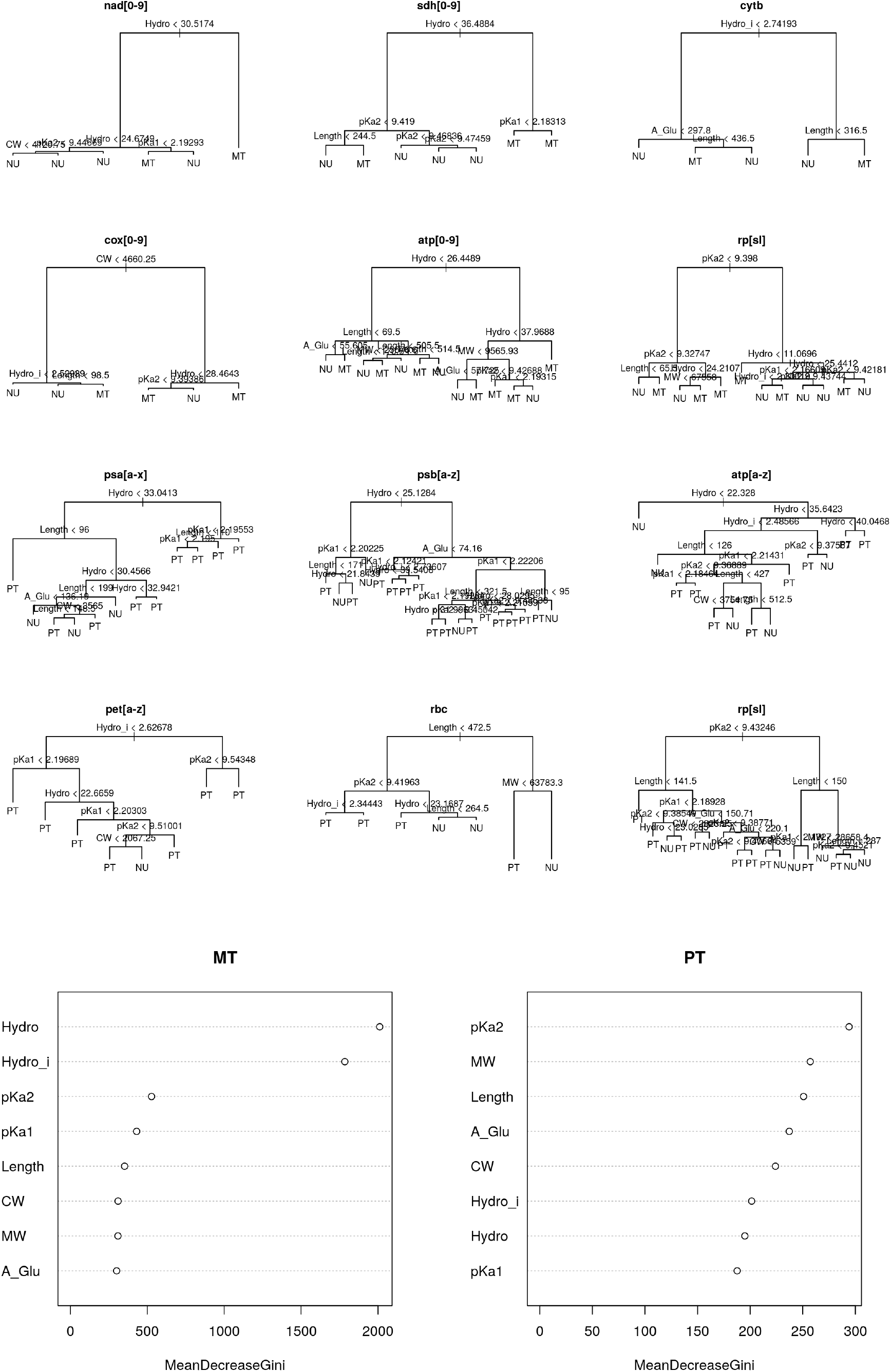
Decision tree and random forest classification for encoding compartment. (top) a set of trees learned to predict encoding compartment for genes in different protein complexes, showing roles for hydrophobicity, pKa, and production energy (CW) as predictive features. (bottom) variance improvement plots for random forest regression for compartment classification across all genes, illustrating the importance of each feature in the predictive outcome. Complexes are labelled with regular expressions describing their gene labels.

